# Plasma cell differentiation, antibody quality, and initial germinal center B cell population depend on glucose influx rate

**DOI:** 10.1101/2023.09.13.557599

**Authors:** Shawna K. Brookens, Sung Hoon Cho, Yeeun Paik, Kaylor Meyer, Ariel L. Raybuck, Chloe Park, Dalton L. Greenwood, Jeffrey C. Rathmell, Mark R. Boothby

## Abstract

Antibody secretion into sera, selection for higher affinity BCR, and the generation of higher Ab affinities are important elements of immune response optimization, and a core function of germinal center reactions. B cell proliferation requires nutrients to support the anabolism inherent in clonal expansion. Glucose usage by GC B cells has been reported to contribute little to their energy needs, with questions raised as to whether or not glucose uptake or glycolysis increases in GC B cells compared to their naïve precursors. Indeed, metabolism can be highly flexible, such that supply shortage along one pathway may be compensated by increased flux on others. We now show that elimination of the glucose transporter GLUT1 after establishment of a pre-immune B cell repertoire, even after initiation of the GC B cell gene expression program, decreased initial GC B cell population numbers, affinity maturation, and PC outputs. Glucose oxidation was heightened in GC B cells, but this hexose flowed more into the pentose phosphate pathway (PPP), whose activity was important in controlling reactive oxygen (ROS) and ASC production. In modeling how glucose usage by B cells promotes the Ab response, the control of ROS appeared insufficient. Surprisingly, the combination of galactose, which mitigated ROS, with provision of mannose - an efficient precursor to glycosylation - supported robust production of and normal Ab secretion by ASC under glucose-free conditions. Collectively, the findings indicate that GC depend on normal glucose influx, especially in PC production, but reveal an unexpected metabolic flexibility in hexose requirements.

**KEY POINTS:**

1. Glucose influx is critical for GC homeostasis, affinity maturation and the generation of Ab-secreting cells.
2. Plasma cell development uses the Pentose Phosphate Pathway, and hexose sugars maintain redox homeostasis.
3. PCs can develop and achieve robust Ab secretion in the absence of glucose using a combination of hexose alternatives.

## INTRODUCTION

The B lymphocyte lineage is integral for effective short and long-lasting humoral immunity. B cells are precursors to antibody secreting cells (ASCs) and memory cells which arise by both germinal center (GC)-dependent and -independent pathways. Optimal antibody responses require B lymphoid cell proliferation and differentiation into memory B and plasma cells, accompanied by diversification of the pre-immune antigen receptor pool. Over recent years, increasing evidence provides some insights into the metabolic demands and reprogramming of B lineage cells during activation, differentiation, and persistence (1, 2). Metabolic requirements evolve as B lineage cells mature and differentiate to support their various functional demands. Uptake of glucose, fatty acids, and amino acids increases during B cell activation and entry into germinal center (GC) reactions (3–7) The usage of glucose by GC B cells is reported to contribute little to their energy needs (8), with questions raised as to whether or not glucose uptake or glycolysis increases in GC B cells compared to their naïve precursors. In contrast, long-lived plasma cells (PC, LLPC) are reported to have increased intake of the glucose analogue 2-(N-Nitrobenz-2-oxa-1,3-diazol-4-yl)amino-2-deoxyglucose (2-NBDG) compared to their short-lived counterparts, and it has been suggested that most glucose consumption by LLPC is used for the glycosylation of antibodies (9, 10). Once taken up into the cell, glucose has multiple potential fates, including usage for glycolysis, the pentose phosphate pathway (PPP), and glycosylation. However, the contributions of glucose to GC biology and plasma cells is still unclear, as are the predominant as well as required pathways for its use throughout the lifespan of mature B linage cells and their progeny.

Glucose enters cells through one of several GLUT transporters expressed on the B cell membrane. Previous groups have provided evidence for the importance of glucose and glycolysis in developing B cells (11–13). A major glucose transporter on lymphocytes, GLUT1 (encoded by *Slc2a1*), was critical for the production of mature B cells and a subsequent Ab response to a hapten (12). Due to the developmental effect, however, it is not clearly established that GLUT1 is needed on B cells after their development into a normal pre-immune repertoire. If so, what steps of cellular differentiation are affected by its absence, and which portions of intermediary metabolism account for the alterations, also are not known.

Accordingly, we sought to elucidate cellular and molecular mechanisms that promote B cell function downstream of glucose and its uptake. To do so, B cell type-specific gene disruption of the glucose transporter GLUT1 was induced in mature mice with a normal pre-immune repertoire for *in vivo* and *in vitro* analyses combined with analyses exploiting *in vitro* glucose starvation. Based on data showing that the oxidative PPP consumes several-fold more glucose than is oxidized upon mitochondrial entry, the impact of inhibiting the PPP on plasma cell differentiation *in vitro* was tested. Limiting glucose transport impaired GCB numbers early in a response or when GLUT1-deficient B cells competed with normal B cells, and quantities of ASC output, Ab concentrations in sera, and affinity maturation required fully intact glucose import in secondary responses.

Because our data and earlier findings (9) suggested that PC have increased demand for glucose compared to GC B cells, we further explored questions about the extent to which glucose is unique in providing for the differentiation and survival of activated B cells as they develop into ASC, and the production of Ab. Redox homeostasis in the B lymphocyte lineage is critical as activated B cells increase intracellular reactive oxygen species upon activation. Moreover, PC differentiation involves enhancing and reshaping antioxidant responses comprised of several partially redundant redox pathways (14, 15). Among the hexoses or simple carbohydrates that could provide fuel and carbons for lymphocyte anabolism, glucose has received most of the attention (16). Nonetheless, several hexoses are available to the metabolic processes in immune cells (17), and simple sugars overlap in their ability to fuel glycolysis, glycosylation, PPP, and glycogen synthesis. Which hexose(s) might suffice to support the capacity of the B lymphocyte lineage to yield secreted Ab remains unclear. To explore this issue, we supplemented B cells with galactose and / or mannose to uncover whether glucose is unique in fueling B cells or supporting antibody production.

## MATERIALS & METHODS

### Mice and immunizations

All animal protocols - reviewed and approved by Vanderbilt University Institutional Animal Care and Use Committee - complied with the National Institutes of Health guidelines for the Care and Use of Experimental Animals. Mice were housed in specified pathogen-free conditions. Both male and female mice, aged 6-10 weeks, were used; sex-specific subgroup analyses did not reveal any significant differences and yielded the same conclusions. To enable tamoxifen-induced, B cell type-specific inactivation of the GLUT1 coding potential, *Slc2a1^f/f^* mice (18) were crossed with *huCD20*-CreER^T2^ (19, 20) or *S1pr2*-CreER ^T2^ (21) transgenic mice; for confirmatory in vitro work with purified B cells, some experiments used *Rosa26*-CreER^T2^ [as in (5, 9, 22)]. Tamoxifen was administered as reported previously (22). To control for potential Cre toxicity in B cell responses or GC B cells (23, 24), age-matched *Slc2a1^+/+^ huCD20*-CreER^T2^ mice were co-housed with *Slc2a1^f/f^ huCD20*-CreER^T2^ mice and used as wild-type controls.

Mice were immunized, or immunized and boosted, by one or two intraperitoneal (i.p.) injections of 100 µg of 4-hydroxy-3-nitrophenylacetyl hapten (NP) conjugated to ovalbumin (NP-ova, Biosearch Technologies, Novato, CA), emulsified in 100 µL Imject^®^ alum (Thermo Fisher Scientific, Pittsburgh, PA), as described previously (22). For mucosal challenges to elicit an anti-carrier Ab response, ovalbumin (50 µg dissolved in 20 µL PBS) was instilled intranasally (i.n.) once daily across seven consecutive days starting three weeks after i.p initial sensitization of mice with NP-ova. Mice were harvested 12 h after the final inhalation. To analyze proliferation rates in vivo with BrdU, or purify GC-phenotype B cells for assays of metabolism or quantitation of recombination at the *Slc2a1* locus, SRBC immunizations were performed as described previously (5, 20), followed by preparative sorting as indicated.

For analyses of cell cycle rates in vivo, mice received intravenous injections of BrdU (100 mg/kg in sterile PBS) (Sigma-Aldrich) 16 and 4 h before harvesting splenocytes. Single-cell suspensions, prepared as described above, were stained for surface markers (B220, GL7, CD95, ** *Sung Hoon!!?!* **) followed by cell fixation, permeabilization, and staining BrdU-containing DNA using the BD APC BrdU Flow Kit (BD-Pharmingen, San Jose, CA) according to manufacturer’s protocol.

For cell transfer experiments, B cells from *S1pr2*-CreER^T2^ or *Slc2a1^f/f^; S1pr2*-CreER^T2^ were purified by depleting CD90^+^ T cells and CD138^+^ cells using biotinylated anti-Thy1.2 Ab and biotinylated anti-CD138 Ab followed by streptavidin-conjugated microbeads (iMag^TM^; BD Biosciences, San Jose CA). B cells from CD45.1; IgH^[a]^ mice were separately purified by the same negative selection. WT CD4^+^ T cells and OT-II CD4^+^ T cells were purified by positive selection with L3T4 anti-CD4 microbeads (Miltyeni Biotech, Auburn CA). *S1pr2*-CreER^T2^ or *Slc2a1^f/f^; S1pr2*-CreER^T2^ B cells (5 × 10^6^ cells per recipient) were mixed with CD45.1; IgH^[a]^ B cells (5 × 10^6^ cells per recipient) and CD4^+^ T cells (5 × 10^6^ cells per recipient; polyclonal : OT-II = 4:1), and injected intravenously into irradiated (7 Gy; 2 split dose of 3.5 Gy with overnight interval, followed by 3 day recovery) CD45.1; IgH^[a]^ recipient mice. Mice were then immunized with NP-OVA as above, treated with tamoxifen at days 3, 5, and 7 after primary immunization, and harvested at 14 day.

### Cell cultures

Splenic B cells were purified from mice using negative selection as previously described (22) or by positive selection with anti-mouse B220 nanobeads (Miltyeni Biotec, Auburn CA). To induce plasma cell differentiation, B cells (seeded at 5 x 10^5^ / mL) were cultured with 5 µg/mL LPS (Sigma-Aldrich, St Louis, MO), 10 ng/mL BAFF (AdipoGen, San Diego, CA), 10 ng/mL IL-4 (Peprotech, Rocky Hill, NJ), 5 ng/mL IL-5 (Peprotech), and 50 nM 4-hydroxy-tamoxifen (4-OHT) (Sigma) and re-fed at day 3. When cultured 2 d or less, cells were in the same conditions after plating at 2 x 10^6^ / mL. Alternatively, B cells were activated with 1 µg/mL anti-CD40 (Tonbo, San Diego CA) and cultured as for LPS blasts. Glucose deprivation and hexose supplementation analyses were performed using glucose-free RPMI (Thermo Fisher, Waltham MA) supplemented with 10% dialyzed Gibco FBS (Thermo-Fisher), 100 U/mL penicillin (Invitrogen, Waltham MA), 100 µg/mL streptomycin (Invitrogen), 3 mM L-glutamine (Invitrogen), non-essential amino acids (NEAA, Invitrogen), 10 mM HEPES (Invitrogen), 0.1 mM 2-mercaptoethanol (Sigma). Where indicated, glucose-free cultures were hexose-supplemented using D-glucose (2 g/L) (Fisher, D-galactose (2 g/L) (Sigma-Aldrich), or D-mannose (125 µM) (Sigma-Aldrich).

To analyze proliferation in vitro, B cells (2 x 10^6^, purified as above) were labeled with 5 µM CellTrace Violet (CTV) (Invitrogen, Waltham, MA) and then activated and cultured as above. Alternatively, rates of S-phase entry were determined by BrdU incorporation, for which day 2 LPS, BAFF, IL-4, and IL-5 cultured cells were pulsed (4 hr) with BrdU (10 µM) (Sigma-Aldrich) prior to flow staining with an anti-BrdU-Alexa fluor 647-conjugated antibody (BD Pharminogen, San Jose CA)], as described (26). Where indicated, oxidative pentose phosphate pathway (PPP) inhibitors 6-aminonicotinamide (6-AN) and dehydroepiandrosterone (DHEA) (Cayman Chemicals, Ann Arbor MI) were added to cultures at the time of activation.

### Real time PCR

RNA was isolated from day 2 LPS blasts seeded with sorted B cells using TRIzol reagent following the manufacturer’s instructions (Life Technologies, Carlsbad, CA). Genomic DNA was extracted from flow-purified naïve and GC B cells using lysis buffer (0.2% SDS, 100 mM Tris base, 5 mM EDTA, and 20 mM NaCl) and proteinase K (20 µg/mL). *Slc2a1*-sequences (+ and fl alleles) were amplified from genomic DNA pools with the primer pair F - CTGTGAGTTCCTGAGACCCTG; R – CCCAGGCAAGGAAGTAGTTC, while the *Slc2a1*^Δ^ allele was amplified with F – CTGTGAGTTCCTGAGACCCTG; R – GACCACGTCTGATGCCAGT. *Slc2a1*^f/f^ ‘locus retention’ (1 - % deleted) was calculated relative to β-actin using the 2-ΔΔCT method and normalized to the fl allele signal. cDNA templates, synthesized using an AMV Reverse Transcriptase kit (Promega, Madison, WI), were amplified using SYBRGreen Power UP Master Mix (Thermo Fisher Scientific) and the following primers for *Slc2a1*-encoded sequences: F-TTCACTGTGGTGTCGCTGTTTG; R-GCTCGGCCACAATGAACCAT. *Slc2a1*–encoded mRNA levels were calculated relative to β-actin using the 2-ΔΔCT method, followed by normalizing experimental samples to the value for wild-type cDNA.

### Flow cytometry

Fluor-conjugated mAbs were purchased from BD Pharmingen (San Jose, CA), Life Technologies, eBiosciences (San Diego, CA), or Tonbo Biosciences. For 2-NBDG uptake analyses, 3 x 10^6^ cells were incubated in PBS (1 h at 37 C) to deplete intracellular glucose stores, stained with Ghost-510 and 2-(N-Nitrobenz-2-oxa-1,3-diazol-4-yl)amino)-2-deoxyglucose (2-NBDG; 60 µM) (Thermo Fisher Scientific) (30 min at 37 C), followed by staining with anti-B220, -IgD, -CD138, -TACI, -NP, -GL7, and -CD38. For detection of GC- and memory-phenotype B cells in the spleens of immunized mice, samples were stained as previously described (22). In brief, 3 x 10^6^ splenocytes were stained with anti-B220, -GL7, -Fas, -IgD, -CD38, NP-APC and a dump channel containing anti-CD11b, -CD11c, -F4/80, -Gr-1, and viability marker (7-AAD or Ghost-Brilliant Violet 510), in selected instances with FITC-anti-human GLUT1 (Alomone Labs, Jerusalem, Israel) in 1% BSA and 0.05% sodium azide in PBS. For detection of PC or activation markers in products of *in vitro* cultures, viable cells (gated via FSC, SSC, and Ghost e450) were stained with fluorophore-conjugated anti-CD138, -B220, -CD69, -MHCII, -CD86, -CD80, or –CD98. For flow analyses of mitochondrial and total intracellular ROS, cells (1-3 x 10^6^) were washed in PBS and stained with 5 µM MitoSOX or 1.25 µM H_2_DCFDA, respectively, along with Ghost-780 in PBS (20 min at 37°C), then washed again (1% BSA in PBS) and further stained with anti-B220, anti-IgD, anti-GL7, -CD138, or -CD38. Samples were analyzed using a FACS Canto flow cytometer driven by BD FACS Diva software or as part of preparative flow purification with a FACS Aria flow sorter. Data were processed using Flow-Jo software (FlowJo LLC, Ashland, OR).

### ELISA and ELIspot

Relative NP-specific Ab concentrations in sera, and secreted Ab in culture supernatants, were measured using capture ELISA as described previously (22). All- or high-affinity hapten-specific Ab were measured using plates coated with NP_20_-BSA or NP_2_-PSA (Biosearch Technologies), respectively. Ovalbumin-specific antibodies were measured using serial dilutions of sera incubated on ovalbumin (1 µg/well)-coated 96-well plates (16 h at 4° C). Total secreted Ab were captured using an anti-Ig(H+L) reagent. Captured Ab retained after rinsing were detected in isotype-specific manner using HRP-conjugated anti-IgM, -IgG1, IgG2c, or -IgE incubation and colorimetric development with Ultra TMB Substrate (Thermo Fisher Scientific).

The frequencies of antibody secreting cells (ASCs) in the bone marrow, lung, or spleen after immunization were measured as described previously (22). For the detection of IgM or IgG1 ASC in day 5 *in vitro* cultures, high protein binding plates (Corning Life Science, Corning NY) were coated with anti-Ig(H+L) (Biosearch Technologies) prior to culture [20 hr, with 100, 200, and 500 cells per well (IgM) or 1000, 2000, and 5000 cells/well (IgG1) and subsequent development with biotinylated anti-IgM or -IgG1 antibodies (Southern Biotechnologies, Birmingham AL). ASC numbers and spot sizes were quantified using an ImmunoSpot Analyzer (Cellular Technology, Shaker Heights, OH) and the densities with readily resolved spots (IgM, 200; IgG1, 2000 cells/well, respectively).

### Radiotracer and glucose uptake and consumption assays

For glucose uptake assays, equal numbers of cells were incubated at 37°C for 15 minutes in glucose uptake buffer (8.1 mM Na_2_HPO_4_, 1.4 mM KH_2_PO_4_, 2.6 mM KCl, 136 mM NaCl, 0.5 mM MgCl_2_, 0.9 mM CaCl_2_, pH 7.4) to deplete intracellular glucose stores as in (4). Samples were incubated in triplicate (10^6^ per sample) with 1 µCi of 2-[1,2-^3^H]-deoxyglucose (20 Ci/mmol; PerkinElmer) in uptake buffer for 4 min at room temperature and immediately spun through a layer of bromo-dodecane into a layer of 20% perchloric acid/8% sucrose to stop the reaction and separate cells from unincorporated 2-[^3^H]-deoxyglucose. Recovered cell lysates were counted by liquid scintillation after this separation from the supernatant. PPP activity and glucose oxidation rates were measured by radiotracer assays using determinations of the amounts of ^14^CO_2_ captured on moist KOH-saturated filter paper after its generation during cultures of cells in 1-[^14^C]-D-glucose or 6-[^14^C]-D-glucose, respectively (4, 12, 27, 28). In brief, PPP activity was assayed using [1-^14^C]-glucose (55 mCi/mmol; American Radiolabeled Chemicals, Inc. St Louis, MO), and glucose oxidation by using [6-^14^C]-glucose (55 mCi/mmol; American Radiolabeled Chemicals, Inc.). Flow-purified GC B cells (B220^+^ IgD^−^ GL7^+^) ex vivo (1 × 10^6^ cells/ml; sample with technical duplication) or naïve B cells (B220^+^ IgD^−^) from SRBC-immunized C57BL/6J mice were incubated in glucose-free RPMI1640 media for 30 min to deplete internal glucose, followed by addition of 0.5 *μ*Ci [^14^C]-glucose, and placement in rubber-stoppered 20 ml scintillation vials containing a center well with 5% KOH-soaked filter paper. Following incubation (37°C for 4 h), radioactivity of trapped ^14^CO_2_ in the filter paper was measured by liquid scintillation counting. In some experiments, purified splenic B cells were activated with LPS or anti-CD40 and cultured with BAFF and IL-4 for 2 days for assays as described above, or for 4 days followed by flow purification of the B cells and CD138^+^ population and assay of PPP activity.

For proton nuclear magnetic resonance spectroscopy (^1^H-NMRS), conditioned culture media were collected after 48 hours of mitogenic activation with and culture in LPS and BAFF and processed as previously reported (29, 30). Briefly, for quantification of metabolites from conditioned supernatant, a total of 50 µL deuterated water (D_2_O) and 50 µL of 0.75% sodium 3-trimethylsilyl-2,2,3,3-tetradeuteropropionate (TSP) in D_2_O were added to 500 µL media and transferred to 5-mm NMR tubes (Wilmad-LabGlass, Kingsport, TN). ^1^H-NMRS spectra were acquired on an Avance III 600 MHz spectrometer equipped with a Triple Resonance CryoProbe (TCI) (Bruker) at 298 K with 7500-Hz spectral width, 32,768 time domain points, 32 scans, and a relaxation delay of 2.7 seconds. The water resonance was suppressed by a gated irradiation centered on the water frequency. The spectra were phased, baseline corrected, and referenced to TSP using Chenomx NMR Suite. Spectral assignments were based on literature values. Glucose consumption was calculated from the difference between the measured concentrations in the media before and after the lymphoblast cultures (30).

### Metabolic flux analyses

Oxygen consumption rate (OCR) and extracellular acidification rate (ECAR) were measured using a Seahorse Bioscience XFe96 extracellular flux analyzer (Agilent Technologies, Santa Clara CA). Activated B cells (2.5 x 10^5^) two day LPS, BAFF, IL-4, and IL-5 were seeded per well of a Cell-Tak (5 µg/mL; Corning) coated plate. Glycolytic and mitochondrial stress tests were performed as previously described (20). Maximum respiration, spare respiratory capacity, and glycolytic reserve were calculated using formulas derived from the software accompanying the Agilent Seahorse platform.

## RESULTS

### Reduced proliferation and plasma cell differentiation of B cells when glucose supply is insufficient

B cells increase glucose uptake upon activation and entry into germinal centers (5, 6, 31). The relative utilization of glucose in different mature B cell subsets beyond the germinal center is less characterized. To approximate the proportional glucose uptake of B cells at different stages after activation, splenocytes were stained *ex vivo* with fluorescently labeled glucose analog 2-(N-Nitrobenz-2-oxa-1,3-diazol-4-yl)amino)-2-deoxyglucose (2-NBDG) two weeks after immunization with the haptenated carrier protein NP-ova (Fig 1A). Flow cytometric analyses of Ag-specific germinal center B cells (GCB), here defined as IgD^−^ GL7^+^, corroborate previous findings reporting ∼2-fold increases in 2-NBDG uptake in GCB compared to the naïve IgD^+^ population (5, 6, 31). Moreover, the Ag-specific population of memory-phenotype B cells, defined here as IgD^−^CD38^hi^, exhibited higher 2-NBDG fluorescence than both the naïve and GCB cell populations. Consistent with earlier data (9, 31), the Ab-secreting cell (ASC) subset had the most avid uptake, demonstrating almost 10-fold higher signal than the naïve population. Recent work has noted that 2-NBDG signals do not necessarily reflect actual glucose uptake (32–34), presumably due to characteristics of the bulky fluorophore coupled to 2-deoxy-D-glucose (2-DG). Radiotracer assays of 2-DG uptake showed that flow-purified GC-phenotype B cells had 3-fold greater influx of glucose compared to naïve controls, whereas this difference was ∼30-fold for *in vitro* B lymphoblasts (Fig. 1B, C). Thus, under the conditions used here, 2-NBDG uptake trends correlated (but mildly underestimated) actual glucose uptake and GC B cells definitely have increased uptake of this metabolically processed hexose.

**Figure 1.**
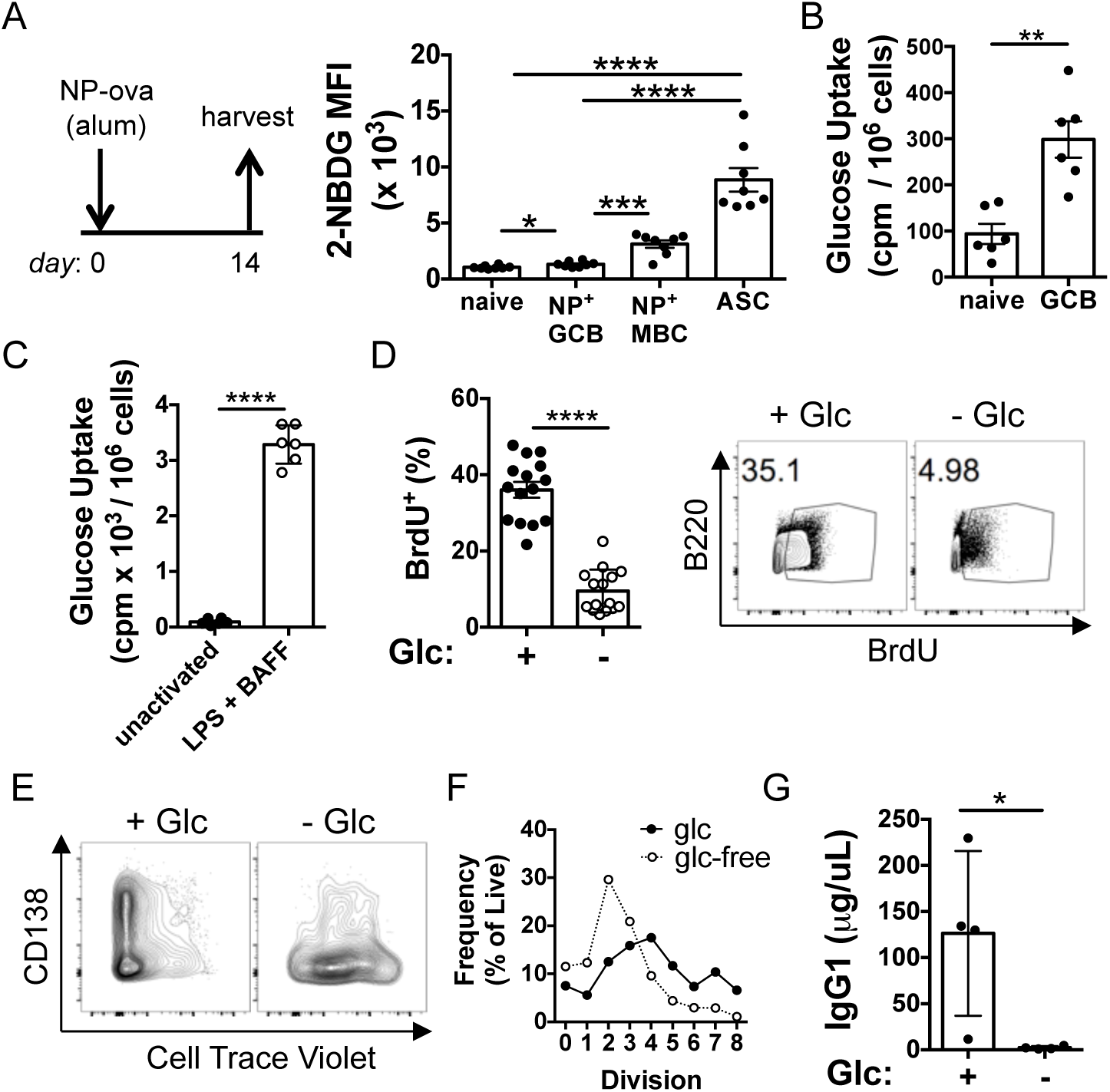
Activation and developmental stage-dependent differences in requisite glucose uptake. (A) Left, schematic depicting immunization with NP-ova in alum to allow measurements of multiple B lineage cell types within the same mouse. Right panel, *ex vivo* 2-NBDG signal in splenic populations: naïve B cells (B220^+^ IgD^+^ GL7^neg^), antigen-specific germinal center B cells (GCB; B220^+^ GL7^+^ IgD^neg^ NP^+^), antigen-specific memory B cells (MBC; B220^+^ CD38^+^ IgD^−^ NP^+^) and antibody secreting cells (ASC; CD138^+^). Each dot shows the mean fluorescence intensity in the indicated cell-type gate, with mean (±SEM) values among the samples (n=8) presented as bar graph. (B) Glucose uptake into naïve (IgD^+^, GL7^neg^, CD138^neg^) and germinal center B cells (IgD^neg^, GL7^+^, CD138^neg^) purified by flow cytometry one week after SRBC immunization, measured using 2-[1,2-^3^H]-deoxyglucose. Each dot shows the measured counts, with mean (±SEM) values among the samples (n=6) as in (A). (C) Glucose uptake into *in vitro*-generated B lymphoblasts, measured using 2-[1,2-^3^H]-deoxyglucose 2 days after activation with LPS and BAFF. Data are shown as in (B). (D) Left panel, BrdU incorporation into DNA of viable cells after culture (2 d) of B lymphoblasts activated with LPS and grown in the presence or absence of glucose (10 mM) in a glucose-free medium supplemented with dialyzed FBS (10%), BAFF, IL-4, and IL-5. Right panel, representative flow cytometric analysis of BrdU incorporation. Data represent four independent experiments totaling 15 B cell preparations from separate mice. (E) Cell Trace Violet (CTV) partitioning four days after activation with LPS and culture in BAFF, IL-4 and IL-5 in the presence or absence of glucose as indicated. (F) ASC differentiation (CD138 expression) as a function of CTV partitioning four days after activation and culture as in (E). (G) IgG1 concentration in the supernatants of these 4 day cultures. * p < 0.05, ** p < 0.01, *** p < 0.001, **** p < 0.0001. Mann-Whitney U test.

Prior work indicated that there was no substantial defect of proliferation or differentiation into ASC when (35) B cells were activated and cultured in glucose-deficient media supplemented with FBS (which contains ∼3 mM glucose, so that glucose is about 300 µM in cultures with 10% FBS). We sought to test the hypothesis that glucose is essential for these B cell processes by supplementing glucose-deficient media with glucose-free dialyzed FBS. Consistent with this model, we found that BrdU incorporation into mitogen-stimulated B cells in glucose-free medium was ∼1/3^rd^ the fraction of BrdU incorporation in parallel samples to which 10 mM glucose was added back (Fig. 1D). Similarly, measurements of Cell Trace Violet (CTV) partitioning between daughter cells showed that B cell division and differentiation to CD138^+^ cells were each drastically reduced in glucose-depleted conditions (Fig 1E, F). These defects culminated in a failure to produce isotype-switched IgG1 Ab *in vitro* (Fig. 1G). In contrast to the decrease in proliferation and differentiation capacity, cell surface expression of B cell activation markers, CD69, MHCII, and CD86 were normal in the absence of glucose (Supplemental Fig. 1A). Although CD80 did exhibit a modest blunting of full induction without glucose, it substantially increased over the 40h period. Interestingly, the amino acid transport component CD98 was increased more on B cells activated in glucose-deficient conditions than on those in glucose-sufficient media, suggesting a compensatory pathway involving increased utilization of amino acids when glucose is unavailable (Supplemental Fig. 1B). We conclude that B cells respond to early activation signals in the absence of glucose, but they require a low level of this hexose for proliferation and differentiation.

### Functional consequences of reduced glucose transport into B cells via *Slc2a1* inactivation

Glucose is imported into B cells through various transporters that include GLUT1, which is encoded by *Slc2a1*. *Cd19*-Cre-driven deletion within *Slc2a1*, a promoter active during B cell development when glucose and glycolysis are critical (11, 13), resulted in 1/3^rd^ as many steady-state splenic B cells and serum Ab compared to wild-type (12). However, mechanistic bases for this observation were not investigated and the experiments did not detect affinity-matured IgG. To investigate the molecular and cellular requirements for glucose flux specifically in mature B cells, we used conditional forms B cell-restricted depletion of GLUT1 via inactivation of *Slc2a1^f/f^* alleles. The tamoxifen-inducible huCD20-CreER^T2^ transgene is active in mature B cells but not their progenitors or plasma cells (19, 36). Mice with B cell-specific deletion of *Slc2a1^f/f^* will be denoted as *Slc2a1^Δ/Δ^*, albeit interchangeably with earlier designations (Glut1 *^Δ/Δ^*or *Glut1^Δ/Δ^*) (12). Quantitative PCR with purified B cells revealed that recombination and deletion of the segment flanked by loxP sites was substantial (∼93% efficient) in the naïve B cells (Fig. 2A) and slightly more complete (∼97.5%) 2 d after mitogenic activation (Fig. 2B). *Slc2a1-* encoded RNA in both naïve and activated B cells purified from tamoxifen-treated *Slc2a1^Δ/Δ^* mice also was substantially decreased compared to controls (Fig. 2A, B). Interestingly, mRNA levels were reduced less than the almost complete recombination of both of the *Slc2a1* gene alleles in both naïve and activated cells. This unexpected but reproducible observation suggests a phenotypic lag, which can be due to factors such as a long t_1/2_ for the primary transcript or mRNA. Consistent with the reported presence of other GLUT transporters on B cells (10, 11, 37) and the phenotypic lag, glucose uptake was reduced but not eliminated in *Slc2a1^Δ/Δ^* samples (Fig 2C). *In vitro*, mitogen-stimulated *Slc2a1^Δ/Δ^* B cells exhibited less robust divisions than their CreER^T2^ control counterparts (Fig 2D). To test if quantitative differences in the surface levels of GLUT1 could affect differentiation, we used an ectodomain-directed antibody to flow-purify naïve GLUT1^hi^ and GLUT1^lo^ B cells that were *Slc2a1^Δ/Δ^*or *Slc2a1^+/+^*. Of note, normal cells with more transporters yielded more CD138^+^ progeny, and the conditional loss-of-function mutation reduced the frequency of these differentiated cells for both populations (GLUT1^hi^; GLUT1^lo^) (Fig. 2E, F). Moreover, the total and division-specific frequencies of B220^lo^ CD138^+^ cells were reduced in *Slc2a1^Δ/Δ^* samples, indicating a division-independent requirement for fully normal glucose uptake in plasma cell differentiation (Fig 2G, H). Collectively, the data on functional impact of *Slc2a1* gene inactivation and reduced glucose transport indicate that proliferation and progression to a plasma cell phenotype in vitro depend on a normal influx of glucose into B cells.

**Figure 2.**
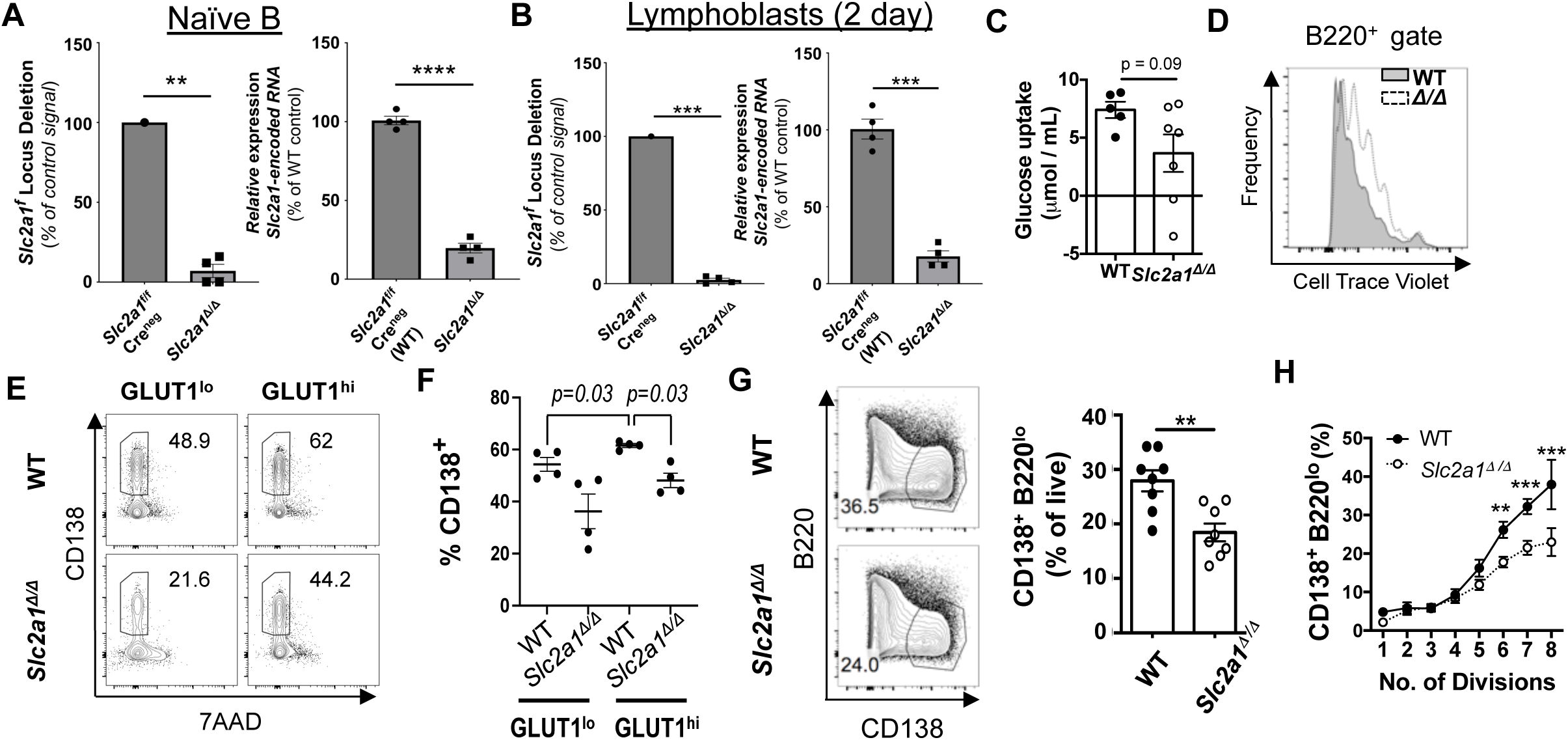
Sufficiency of glucose influx is critical for plasmablast differentiation *in vitro*. (A, B) Deletion efficiency of *Slc2a1* fl alleles in vivo (A) and after activation and culture (2 d) in vitro (B). (A) Naïve-phenotype B cells were flow-purified from spleens of CreER^T2^ mice (*Slc2a1^f/f^*and wild-type) after *in vivo* tamoxifen injections. Shown are both the extent of deletion (bar graph to left, i.e., reduction of the signal for the *fl* allele in qPCR measurements of DNA from *Slc2a1* f/f B cells after tamoxifen injections in CreER^T2^ mice), and the level of *Slc2a1*-encoded RNA (right panel). and culture (2 d) with LPS, BAFF, and 4-hydroxytamoxifen. (B) Flow-purified naïve B cells from experiments in (A) were activated cultured 2 d in BAFF and LPS, followed by DNA and RNA purification and q(RT2)PCR to quantitate *Slc2a1* sequences as in (A). Left and right bar graphs are as in (A). Each shows data from four (4) mice of each genotype (non-recombined / deleted vs deleted) in two biologically independent replicate experiments of 2 vs 2 mice each, with p values calculated by unpaired Student’s t-test. (C) Glucose consumption (extraction from culture media) determined by H^+^ NMR measurements using the supernatants of B lymphoblasts [wild-type vs *Slc2a1^Δ/Δ^*, designated *Glut1^Δ/Δ^* in earlier work (12)] after culture (2 d) with LPS, BAFF, and 4-hydroxytamoxifen. Data represent two independent experiments with wild-type (5) and *Slc2a1^Δ/Δ^*(7) mice. (D) Flow cytometric measurement of CTV partitioning of wild-type and GLUT1-deficient B220^+^ cells 4 d after activation and culture as in (A). (E, F) GLUT1 expression level influences CD138^+^ generation in vitro. Naïve B cells, separated as GLUT1^lo^ versus GLUT1^hi^ using an ectodomain-directed anti-GLUT1, were flow-purified, followed by activation and culture (5 d) in LPS, BAFF, IL-4, and IL-5. (E) Shown are representative flow plots of CD138 staining at the end of cultures, with inset numbers providing the percentages of CD138^+^ events from the WT and *Slc2a1^Δ/Δ^* B cells. (F) Quantified frequencies of CD138^+^ cells after the cultures as in (F), starting from flow-purified B cells of the wild-type (n=4) and B cell-specific *Slc2a11^Δ/Δ^*gene inactivated (n = 4) mice in two independent experiments. p values were calculated by unpaired Student’s t-test. (G) Representative plot (left) and quantification (right) of the frequencies of CD138^+^ B220^lo^ cells after activation and culture as in (D). (H) Shown are the frequencies of CD138^+^ B220^lo^ cells at each cellular division 4 days after activation. Data are aggregated from three independent replicate experiments using wild-type (n = 7) and *Slc2a1^Δ/Δ^* (n = 8) mice. The probabilities that the null hypothesis would be correct were * p < 0.05, ** p < 0.01, *** p < 0.001 by Mann-Whitney U testing.

We next sought to test *in vivo* consequences of acute depletion of GLUT1 from mature B cells on humoral responses. First, mice were immunized with either SRBC or NP-ovalbumin 10 d after starting injections of *Slc2a1^f/f^* huCD20-CreER^T2^ and wild-type control mice (*Slc2a1^+/+^* huCD20-CreER^T2^) with tamoxifen (Fig. 3). After immunizations with SRBC, the frequencies of GC B cells were substantially lower in the *Slc2a1^Δ/Δ^* subjects compared to controls (Fig 3B, C). Although the trend of data was that the fraction of GC B cells in S-phase was reduced (Fig 3D), the difference did not achieve statistical significance – which may in part relate to counter-selection and inclusion of B cells with non-deleted *Slc2a1* in these GC (Fig 3E). Of note, however, the decrease in frequencies of BrdU^+^ events was more significant for activated (IgD^neg^) B cells that were GL7^neg^ – a population that has been treated as “activated precursors” or pre-GC blasts (38). This result shows that proliferation amongst some B cells in vivo is reduced by inactivation of *Slc2a1* at the outset. We used the NP-ovalbumin hapten-carrier system (Fig. 3F), to extend these data in a system that facilitates quantitation of high- and all-affinity ASC and Ab in sera. Of note, the anti-NP repertoire is immunodominant (39–41), allowing enumeration of the Ag-binding B cell population in the early GC response. Mice with acute deletion of *Slc2a1* in mature B cells generated less NP-binding GCB in response to NP-ova at 7 days after immunization, but overall GC B cell prevalence appeared unaffected (Fig. 3G). Of note, the fraction of the recombined *Slc2a1* fl alleles in the flow-purified GC B cell population at the time of harvest was less uniform than in naïve or in vitro-activated B cells (Fig. 3E). Nonetheless, purified NP-binding IgM and IgG1 ASCs in the spleen (Fig. 3H) and circulating anti-NP IgM and IgG1 were markedly diminished in *Slc2a1^f/f^*huCD20-CreER^T2^ mice one week after NP-ova immunization (Fig. 3I).

**Figure 3.**
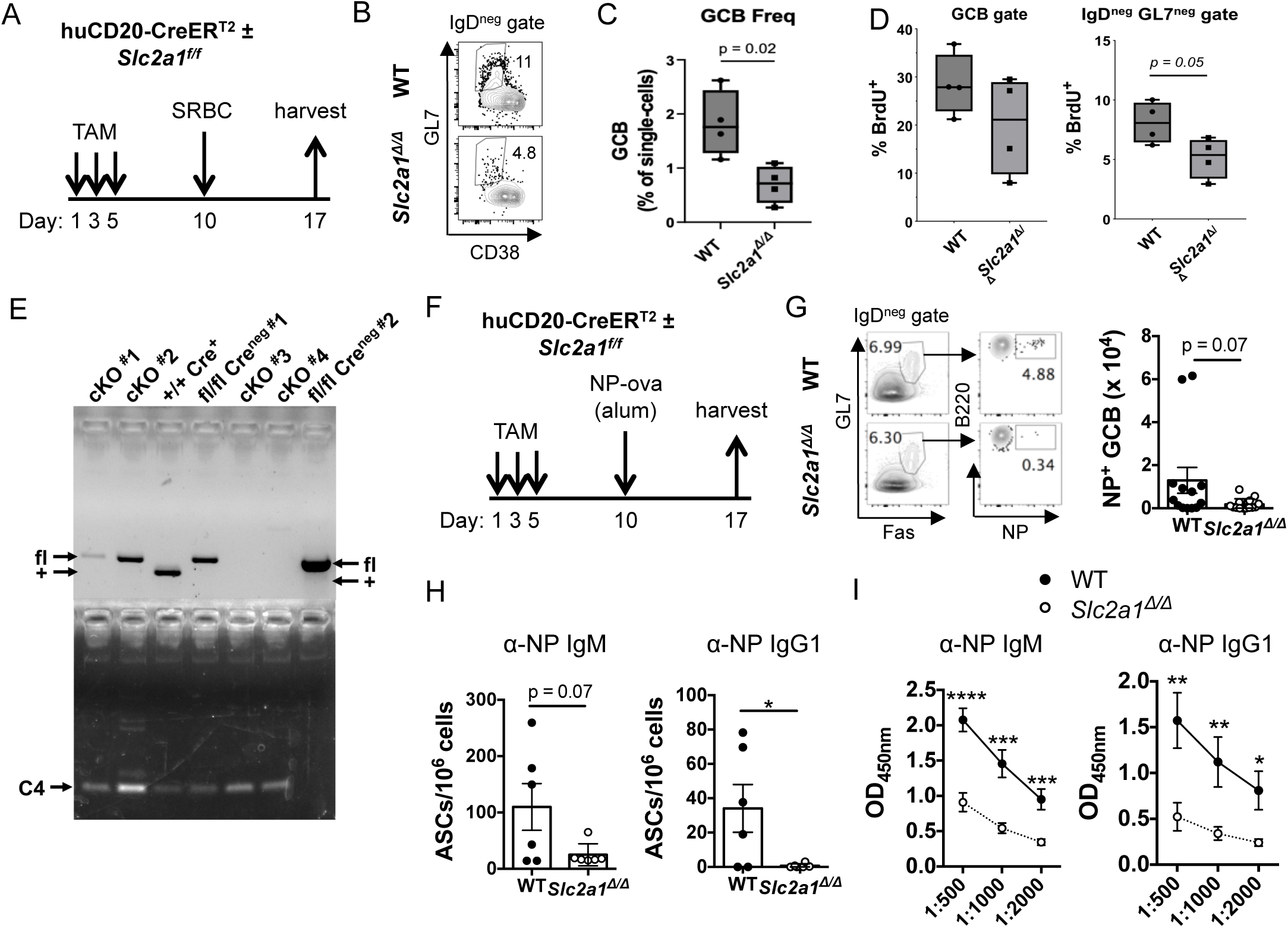
Early immunization-induced germinal center formation, ASC development, and Ab production *in vivo* require normal glucose transport. (A) Schematic depicting immunization with SRBC after inactivation of *Slc2a1* in mature B cells. Mice (huCD20-CreER^T2^, *Slc2a1*^+/+^ or CreER ^T2^, *Slc2a1*^f/f^) were harvested 1 wk after immunization. Representative flow plots of viable splenic IgD^neg^ B cells (B) and aggregate data for two replicate experiments (C) (n=2 for each genotype in each). (D) In vivo BrdU incorporation into IgD^neg^ B cells of the mice in (A-C). Shown are aggregate data, with representative flow plots in Supplemental Fig. 1D. (E) Shown are PCR results for the *Slc2a1* fl allele in flow-purified GC B cells of the mice in (A-C). (F) Schematic depicting immunization with NP-ova after inactivation of *Slc2a1* in mature B cells. Mice (CreER^T2^, *Slc2a1*^+/+^ or CreER ^T2^, *Slc2a1*^f/f^) were harvested 1 wk after immunization. (G) Representative plots (left panel) and total numbers of splenic Ag-binding GCB (right panel) after a single immunization as in (F). Each dot represents an independent mouse of the indicated genotype, with the mean (±SEM) values depicted by bar graph. (H) Ab-secreting cells (ASC) that produced NP-binding IgM and IgG1 one week after immunization were enumerated by ELISpot assays with spleen cell suspensions of the NP-ova-immunized mice as in (G). (I) Relative concentrations of circulating NP-specific IgM and IgG1 in the sera of wild-type and *Slc2a1^Δ/Δ^* mice one week after NP-ova immunization determined by ELISA. Data represent 4 independent experiments with wild-type (n = 13) and *Slc2a1^Δ/Δ^* (n=14) mice. Probabilities of the null hypothesis being correct were * p < 0.05, ** p < 0.01, *** p < 0.001, **** p < 0.0001.

Although GLUT1 loss initiated prior to immunization appeared to reduce the prevalence and numbers of Ag-specific GC B cells, this finding could arise through an effect on B cells prior to entry into the secondary follicle. Moreover, the early ASC outputs and Ab responses include inputs from extrafollicular or recall differentiation of PC (42–45). To test if rates of glucose import into GC B cells affect physiology or outputs from the secondary follicle, we used conditional deletion of *Slc2a1* driven by the *S1pr2*-CreER^T2^ transgene (21, 46). This established model exploits the selectively high (>32-fold increased) expression of the *S1pr2* gene in GC cells (47). Tfh cells also express *S1pr2* mRNA, and global inactivation of the gene is deleterious to the mouse health. Accordingly, we transferred B cells (*S1pr2*-CreER^T2^, *Slc2a1^+/+^* vs *S1pr2*-CreER^T2^, *Slc2a1^f/f^*) into non-lethally irradiated recipients that, relative to the donor genotype, were allotype-disparate at both the IgH and CD45 loci (CD45.1, IgHa) (Fig. 4A). These chimeric mice were immunized and tamoxifen injected after the initiation of GC so that donor and recipient GC B cells (Fig. 4B-D) and Ag-specific ASC (Fig. 4E) could be quantified after harvest. The results showed lower GC B cell population and reduced outputs of ASCs (Fig. 4, B-D, and E, respectively). Thus, GLUT1 reduction on GC B cells compromised their physiology and capacity to generate PC. Collectively, the data indicate that although in some settings the impact on the ASC population may be greater than that on GC B cell homeostasis, the generation or maintenance of GC B cells, ASCs and antibodies in response to immunization require a full capacity to import glucose.

**Figure 4.**
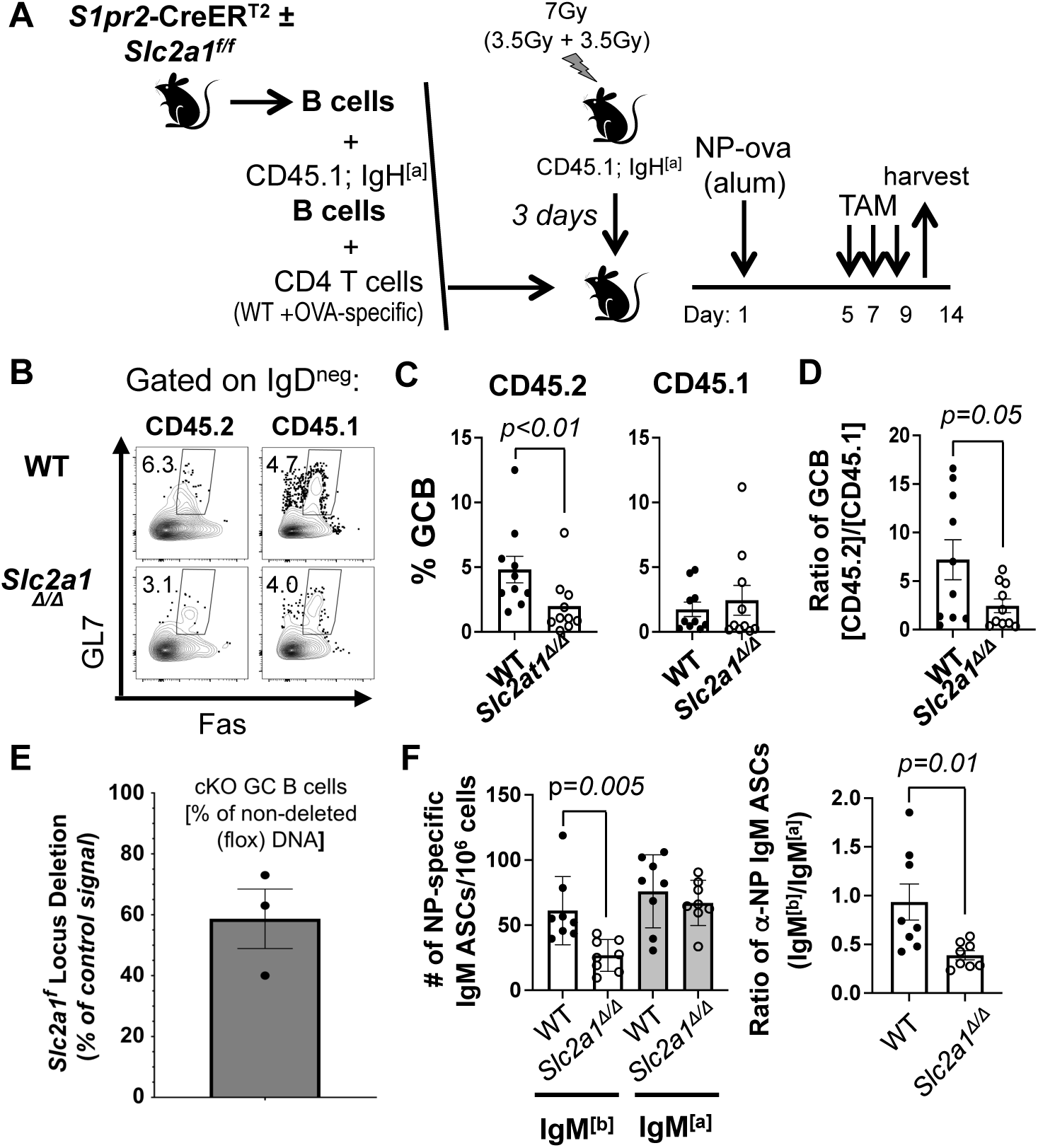
GLUT1 expression within GC B cells regulates their prevalence and PC generation. (A) Schematic model of adoptive transfer and immunization to test the effect of GC B cell-specific role of GLUT1. B cells from *S1pr2*-CreER^T2^ mice (wild-type or *Slc2a1^f/f^*) were mixed with CD45.1; IgH^[a]^ B cells and CD4 T cells, then transferred into irradiated CD45.1; IgH^[a]^ recipient mice followed by immunization with NP-ova. Starting 5 d thereafter, mice were treated with tamoxifen and harvested at day 14. (B-D) Representative flow plots (B), quantified frequencies of Fas^+^ GL7^+^ cells in the B220^+^ IgD^neg^ Dump^neg^ gate (Dump: CD11b, CD11c, F4/80, Gr1, 7-AAD) after gating to distinguish CD45.1 versus CD45.2 (C), and (D) the ratios of CD45.2 GCB / CD45.1 GC B cells. (E) qPCR measurement of *Slc2a1* floxed allele in freshly sorted CD19^+^ IgD^−^ GL7^+^ CD38^−^ GC B cells from tamoxifen-treated WT (*S1pr2*-CreER^T2^) and Glut1 cKO^GCB^ (*Slc2a1^f/f^*; *S1pr2*-CreER^T2^) mice. (F) Numbers of all-affinity NP-specific IgM^[a]^ and IgM^[b]^ ASCs that derived from wild-type (•) and GCB-specific *Slc2a1^Δ/Δ^* (°) B cells (left), and the ratios of NP-specific IgM^[b]^ ASC/ NP-specific IgM^[a]^ ASC harvested from each recipient (right) (n = 8 wild-type *S1pr2*-CreER^T2^ B cell recipients and n = 8 mice that received GC-specific *Slc2a1^Δ/Δ^*B cells in three independent experiments). P values for the likelihood the null hypothesis is correct were calculated by unpaired Student’s t-test.

To investigate longer-term effects of B cell-intrinsic *Slc2a1* on humoral immunity, huCD20-CreER^T2^ mice (*Slc2a1* wild-type versus *Slc2a1^f/f^*) were harvested four weeks after NP-ova immunization and a booster at week three to elicit secondary germinal center responses (Fig 5A). In contrast to the reduced initial population of NP-specific *Slc2a1*-deficient GCB observed one week post immunization (Fig 3B; Fig. 4B-D), *Slc2a1*-deficient GCB were recovered in numbers comparable to wild-type controls in the secondary response (Fig 5B). This result suggests that GLUT1 expression on B cells may differentially influence primary versus secondary GC kinetics or ability to compete and respond to limited antigen, or the ability to form and reactivate memory B cells upon subsequent antigen exposure. Mice harboring *Slc2a1*-deficient B cells exhibited ∼2 fold more NP-specific memory-phenotype splenic B cells compared to wild-type controls in this model of immune exposures (Fig 5C). In sharp contrast, IgM and IgG1 secreting cells that produce antibody with all affinities for NP were modestly reduced in the spleens of *Slc2a1^Δ/Δ^* mice, and high affinity IgG1 secretors were severely ablated (Fig 5D). Consistent with the observed diminution of ASCs, circulating high-affinity anti-NP IgG1 and IgG2c were reduced by the inactivation of *Slc2a1* even when the all-affinity antibody concentrations were comparable (Fig 5E, F). Thus, affinity maturation was reduced for both switched isotypes in the primed and boosted mice whose B lineage cells lacked GLUT1 (Fig. 5G). These data provide evidence that glucose import influences the balance between outputs of memory B cells and ASC, and that its influx is especially critical for the production of high affinity class-switched antibodies that are characteristic outputs of GC reactions.

**Figure 5.**
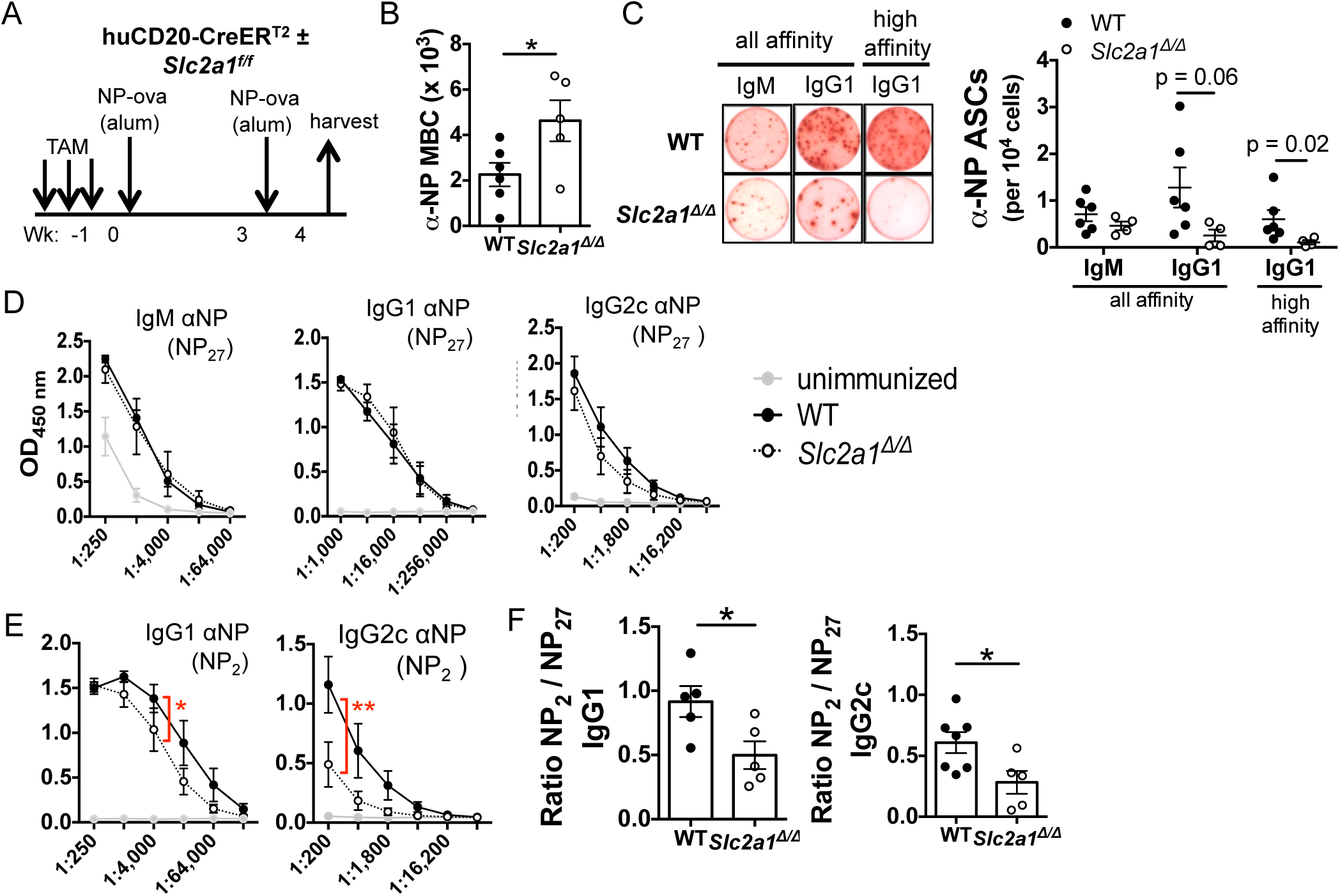
Glucose flux into B lineage cells ASC generation and the production of higher affinity antibodies and restrains early MBC formation. (A) Schematic depicting prime-boost hapten-carrier immunizations after depletion of *Slc2a1* from mature B cells. Mice immunized with NP-ova after tamoxifen-induced conversion of *Slc2a1*from *f/f* to *Δ/Δ* in mature B cells were harvested at week 4, one week after a booster immunization. (B) Total numbers of splenic NP^+^ memory-phenotype (B220^+^ IgD^neg^ GL7^neg^ CD38^++^) B cells at harvest. (C) Left panel, representative ELISpot wells (5 x 10^5^ splenocytes seeded per well) for quantitation of splenic ASCs secreting NP-specific IgM and IgG1 (all- and high-affinity), as indicated. Right panel, splenic ASC prevalence in individual mice, with each dot representing an individual subject. (D) Relative concentrations of all-affinity NP-specific IgM, IgG1, and IgG2c of unimmunized (•), wildtype (•), and *Slc2a1^Δ/Δ^* (°) mice at the time of harvest (1 wk after boost), measured by ELISA across serial four-fold dilutions of the indicated sera. Shown are mean (±SEM) absorbances at each dilution for mice whose B cells were of the indicated genotype. (E) High-affinity NP-specific IgG1 and IgG2c in the sera at the time of harvest, as in (D) but using low-valency NP (NP_2_–PSA). (F) Affinity maturation of IgG1 and IgG2c Ab, calculated as ratios of high-affinity (captured on NP_2_–PSA) to all-affinity (captured on NP_27_–BSA) OD_450_ ELISA values in the linear range (1:16,000 dilution for IgG1 or 1:200 for IgG2c). Data are representative of wild-type (n = 7) and *Slc2a1^Δ/Δ^*(n = 5) mice in two independent experiments. * p < 0.05, ** p < 0.01. Mann-Whitney U test (C), two-way ANOVA (E), Student’s t-test (B, E, F).

We extended the evidence that *Slc2a1* in B cells promotes humoral immunity using an allergic lung inflammation model where mice were primed with NP-ova in the periphery prior to consecutive ovalbumin inhalations to elicit ova-specific B cells in the lung (Supplemental Fig 2A). Akin to previous observations, frequencies of GCB detected in the mediastinal lymph nodes were modestly decreased, accompanied by major defects in antigen-specific IgG1 ASCs in the lungs (anti-ova ASCs) and bone marrows (anti-ova and anti-NP ASCs) of *Slc2a1*-deficient mice (Supplemental Fig 2B-D). The defect of ASC generation caused by interference with GLUT1 expression coincided with attenuated anti-ova IgE and IgG1 circulating antibody (∼0.11x and 0.25x control concentrations, respectively) with modest declines in circulating anti-NP IgM and IgG1 (Supplemental Fig 2E-G). Of note, ELISA quantitation showed that *Slc2a1^Δ/Δ^* B cells supported less affinity maturation, with a reduced ratio of high-to all-affinity anti-NP IgG1 (Supplemental Fig 2H), mirroring evidence that uptake of glucose in B cells promotes affinity maturation (Fig 5G). These data provide new insight into the cellular stages and antibody outputs enhanced by GLUT1-dependent glucose import into B cells. Specifically, normal influx supports GCB formation, affinity maturation, and generation of ASCs in several immunization models.

### Glucose flux in B cells and ASC partitions between pyruvate oxidation and the pentose phosphate pathway (PPP)

To investigate the mechanisms relating to effects of *Slc2a1* inactivation on B cell progression to antibody production, we next measured metabolic fates of glucose in activated B cells. Once imported, glucose is routed to many different pathways in activated B cells to support biosynthetic and energetic demands [(3, 35, 48); reviewed in (2)]. Hexokinase converts glucose to glucose-6-phosphate (G6P), which can then be metabolized via glycolysis to generate intermediates and ultimately pyruvate. Alternatively, G6P can be shunted into the Pentose Phosphate Pathway (PPP), which regenerates NADPH from NADP^+^, to maintain redox balance in B cells (49) and provide reducing equivalents for biosynthetic pathways. The PPP also generates substrates used for endogenous nucleotide and amino acid biosynthesis (50) (Fig 6A). Metabolic flux analyses revealed that activated GLUT1-deficient B cells had substantially lower rates of glucose-stimulated extracellular acidification (ECAR) (Fig 6B), likely reflecting lower flux of glucose through glycolysis to yield pyruvate, and less lactate production. The *Slc2a1^Δ/Δ^* B lymphoblasts had virtually no glycolytic reserve, i.e., no increase in ECAR upon treatment with the mitochondrial ATP synthase inhibitor oligomycin (Fig 6C), indicating that GLUT1 expression supports enhanced glycolytic capacity in metabolically stressful conditions. Previous *in vitro* analyses of activated B cells suggested that glucose is used primarily for biosynthetic pathways with little contribution to oxidative phosphorylation (OXPHOS) (3, 35). However, interference with GLUT1 expression impaired maximal respiration and spare respiratory capacity of activated B cells, suggesting that the fraction of glucose fueling OXPHOS is necessary for optimal mitochondrial function (Fig 6D, E). These data indicate that GLUT1 expression heightens both glycolytic and mitochondrial function during B cell activation.

**Figure 6.**
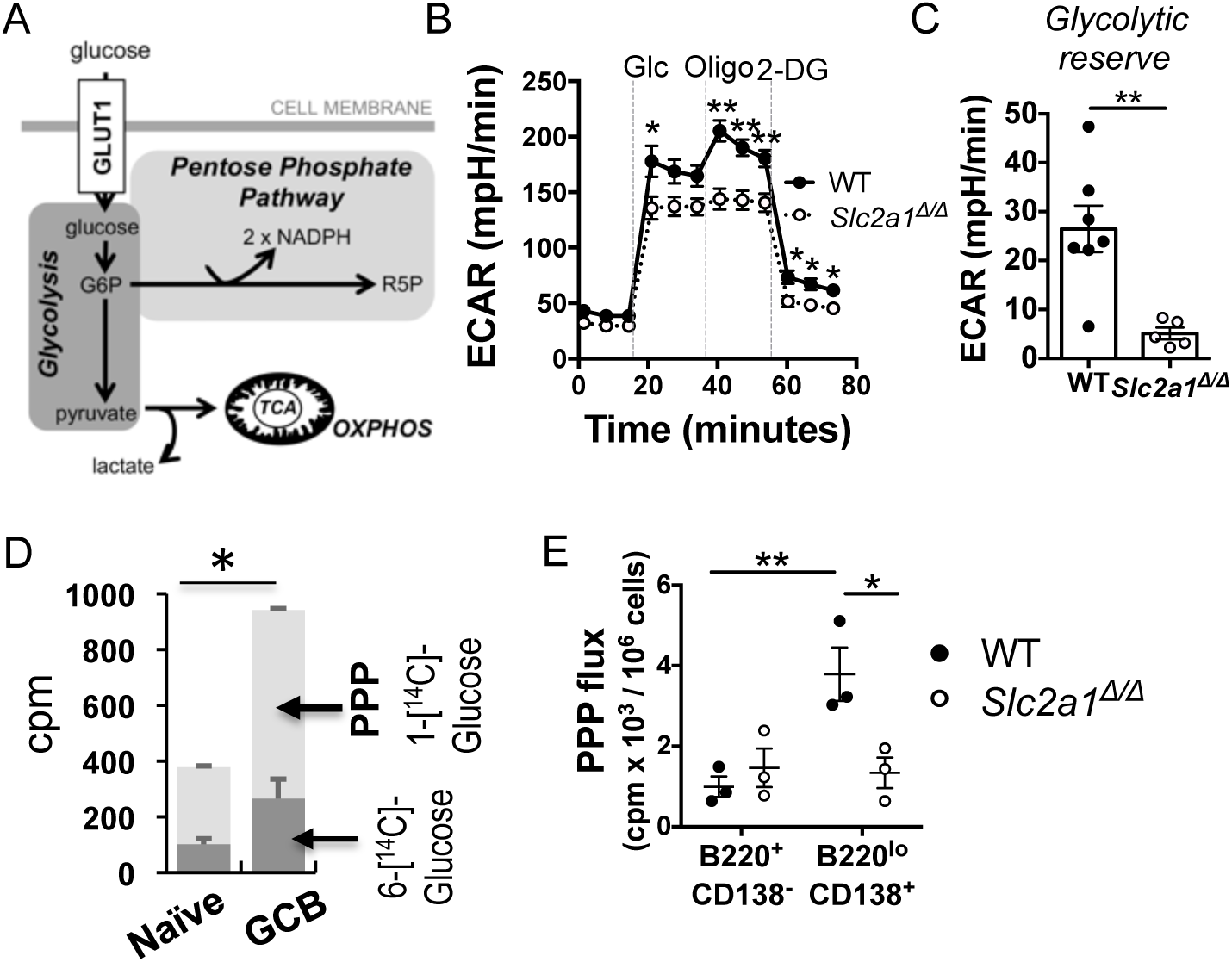
GLUT1 supports metabolic flux in B lymphoblasts and plasma cells. (A) Schematic showing two major fates of glucose-6-phosphate (G6P) generated after glucose entry through the GLUT1 transporter, i.e., glycolysis and shunting into the Pentose Phosphate Pathway. (B) Extracellular acidification rates (ECAR) during metabolic flux analyses conducted as Glycolytic Stress Tests of lymphoblasts. B cells (wildtype or *Slc2a1^Δ/Δ^*) were assayed after activation and culture (2 d) with LPS, BAFF, IL-4, and IL-5. (C) Glycolytic reserve of wild-type and *Slc2a1^Δ/Δ^* B cells calculated from assays in (B). Dots and bar graphing are as in previous figure panels. (D) PPP activity and glucose oxidation determined by 1-[^14^C]-glucose and 6-[^14^C]-glucose conversion to ^14^CO_2_ in the indicated flow-purified splenocyte populations after SRBC immunization. (E) PPP activity determined by 1-[^14^C]-glucose conversion to ^14^CO_2_ after short-term cultures of flow purified wild-type and *Slc2a1^Δ/Δ^*B cells (B220^+^ CD138^neg^) and ASCs (B220^lo^ CD138^+^). Purified B cells were activated and cultured (4 d) with LPS, BAFF, IL-4, and IL-5. Data represent two (B, C) or three (D, E) independent experiments, each with multiple B cell preparations from separate mice. * p < 0.05, ** p < 0.01.

To quantify the oxidation of glucose via glycolysis and the TCA cycle relative to usage by the PPP in B cells, we measured the generation of ^14^CO_2_ from [6-^14^C]- and [1-^14^C]-glucose, respectively (27, 28, 51). Compared to naïve B cells, flow-purified GCB exhibited increases in both [1-^14^C]- and [6-^14^C]-glucose indicating that the increased glucose uptake of GCB feeds into both glycolysis/TCA cycle and PPP (Fig. 6F). Of note, in both naïve and GC B cells, flux of glucose through the PPP activity is about 2-fold higher than glucose oxidation through glycolysis and mitochondrial use of pyruvate to generate acetyl-CoA rather than lactate. *In vitro*-activated B cells had rates of glucose oxidation similar to GCB cells, but the *in vitro* blasts dramatically increased glucose use by the PPP (∼10-fold) compared to *ex vivo* samples (Supplemental Fig. 3A). Glucose uptake for ASC is many-fold greater than for B cells (Fig 1A). The majority of this sugar was diverted to meet demands for glycosylation of secreted Ab when plasma cells were analyzed *ex vivo* in the absence of mannose (9). To determine if GLUT1-dependent glucose uptake in ASC supports flux through the PPP, we measured [6-^14^C]-glucose labeled CO_2_ in flow-purified B cells (B220^+^ CD138^neg^) and differentiated ASC (B220^lo^ CD138^+^) four days after *in vitro* stimulation. PPP activity in undifferentiated B220^+^ cells was unaltered despite the loss of GLUT1 expression (Fig. 6G). By contrast, wild-type ASC substantially increased PPP activity compared to undifferentiated B220^+^ cells in a manner that was almost completely GLUT1-dependent. These data indicate that the increased demand for glucose in ASC was, at least in part, directed to increased PPP flux in these differentiated cells.

The oxidative branch of the PPP is a central player in maintaining redox homeostasis in cells due to the generation of NADPH (50). To determine the effect of GLUT1 loss on oxidative stress in B cells, we measured mitochondrial-derived ROS (mtROS) in splenic naïve B cells, GCB, MBC, and ASC one week after immunization with NP-ova (Fig. 7A). In contrast to B cells, in which GLUT1 loss did not affect mitoSOX signal, this surrogate for mitochondrial reactive oxygen species (mtROS) was 2-fold greater in the *Slc2a1^Δ/Δ^* ASC subset compared to wild-type controls. This finding indicated that sufficient glucose import is critical for maintaining proper mtROS levels specifically in the ASC population. Collectively, the data indicate that glucose import supports glycolytic and mitochondrial functions of B lymphoblasts and is critical for the PPP and redox homeostasis - particularly in differentiated ASC.

**Figure 7.**
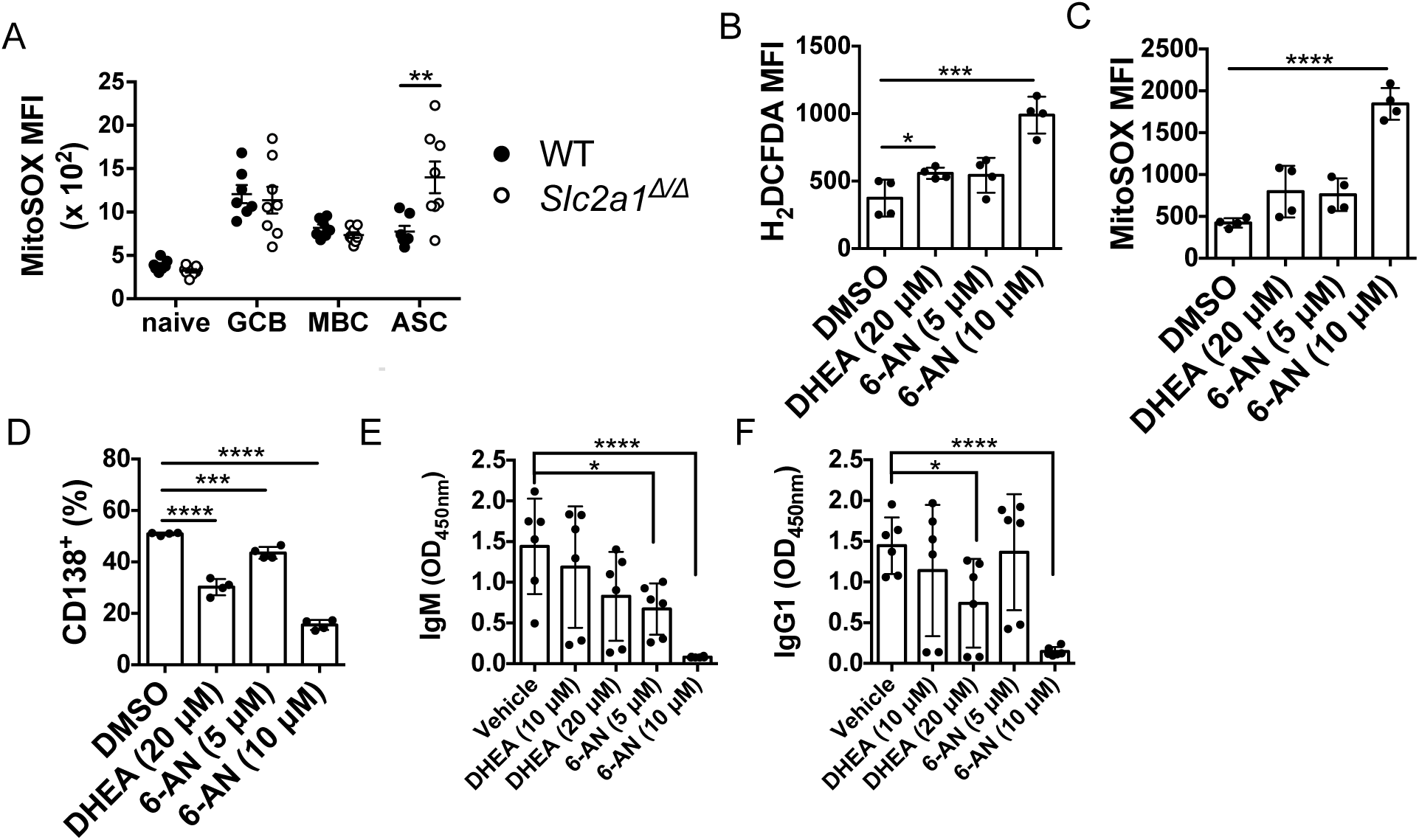
Sufficient activity of pentose phosphate pathway is critical for ROS regulation and the generation of antibody secreting cells. (A) Mitochondrial reactive oxygen species in different B cell subsets measured by ex vivo MitoSOX staining and flow cytometry using mice (wildtype and *Slc2a1^Δ/Δ^*) harvested one week after NP-ova immunization. (B, C) Total cellular ROS (B) and (C) mitochondrial ROS in B lymphoblasts determined by H_2_DCFDA and MitoSOX staining, respectively, followed by flow cytometry. Shown are mean fluorescence intensity (MFI) values for each independent experiment after B cell activation with LPS and culture (2 d) in BAFF, IL-4, and IL-5 in the presence of PPP inhibitors DHEA and 6-AN as indicated. (D) Frequency of CD138^+^ cells after culture ± PPP inhibitor treatment. Cells were activated and cultured as in (B), followed by flow cytometry. (E) Relative concentrations of IgM and (F) IgG1 in supernatants 5 days after B cell activation and culture (5 d) as in (A-D), in the presence of PPP inhibitors as indicated.. Shown are mean (±SEM) ELISA results from three independent experiments and samples, using supernatants at dilutions established as being in the linear range (1:4,000 and 1:1,000 for detection of IgM and IgG1, respectively). * p < 0.05, *** p < 0.001, **** p < 0.0001.

### The PPP promotes ASC development and Ab secretion

In light of these data and the impediment to PC differentiation when glucose influx was restricted, we explored the reliance of ASC development on the PPP. Accordingly, B cells were activated *in vitro* and treated with DHEA, a non-competitive inhibitor glucose-6-phosphate dehydrogenase (G6PD), or 6-AN, an inhibitor of both G6PD and 6-phosphogluconate dehydrogenase (6-PGD) enzymes of the PPP. Activation of B cells in the presence of either DHEA or 6-AN decreased PPP flux (Supplemental Fig. 3B) (52) and increased intracellular ROS and mtROS in lymphoblasts 2 d after mitogen stimulation (Fig 7B, C). Of note, DHEA and 6-AN each also reduced frequencies of differentiated CD138^+^ cells and Ab concentrations in culture supernatants (Fig. 7D-F). These data provide evidence that the capacity to achieve normal flux of glucose through the PPP is critical for redox balance, ASC generation, and antibody production.

### Hexoses mitigate oxidative stress but glucose is best during PC development and function

We next sought to elucidate requirements for and relative contributions of metabolic pathways critical for the generation of ASC and antibody production by culturing cells with different hexoses, which are preferentially metabolized via distinct pathways (53) (Fig. 8A). Substitution of galactose for glucose in cell culture media has been used to decrease glycolytic flux but is thought to preserve PPP activity (54, 55) while mitochondrial respiration can anaplerosis. Accordingly, addition of galactose to hexose-free base medium in extracellular flux assays with activated B cells failed to support increased ECAR (Fig 8B; Supplemental Fig 3C) or respiration (OCR) (Fig. 8C; Supplemental Fig 3D) compared to glucose supplementation. These findings indicate that the G6P generated by the Leloir pathway in activated B cells fails to yield pyruvate, and suggestive of preferential shunting into the PPP. The initial oxidative limb of the PPP results in net reduction of NADP^+^ to NADPH that can be used for resolving ROS. Accordingly, we next tested the relative capacities of glucose versus galactose in promoting redox homeostasis of activated B cells. Consistent with observations with *Slc2a1^Δ/Δ^*deficient ASC, hexose deprivation led to a 3-fold increase in intracellular ROS and mtROS in activated B cells compared to glucose-sufficient conditions (Fig. 8D, E). Notably, addition of galactose in glucose-free conditions restrained steady-state intracellular ROS and mtROS to the same extent as glucose. This finding suggests that galactose can feed the oxidative PPP, with generation of NADPH, to an extent sufficient for redox homeostasis.

**Figure 8.**
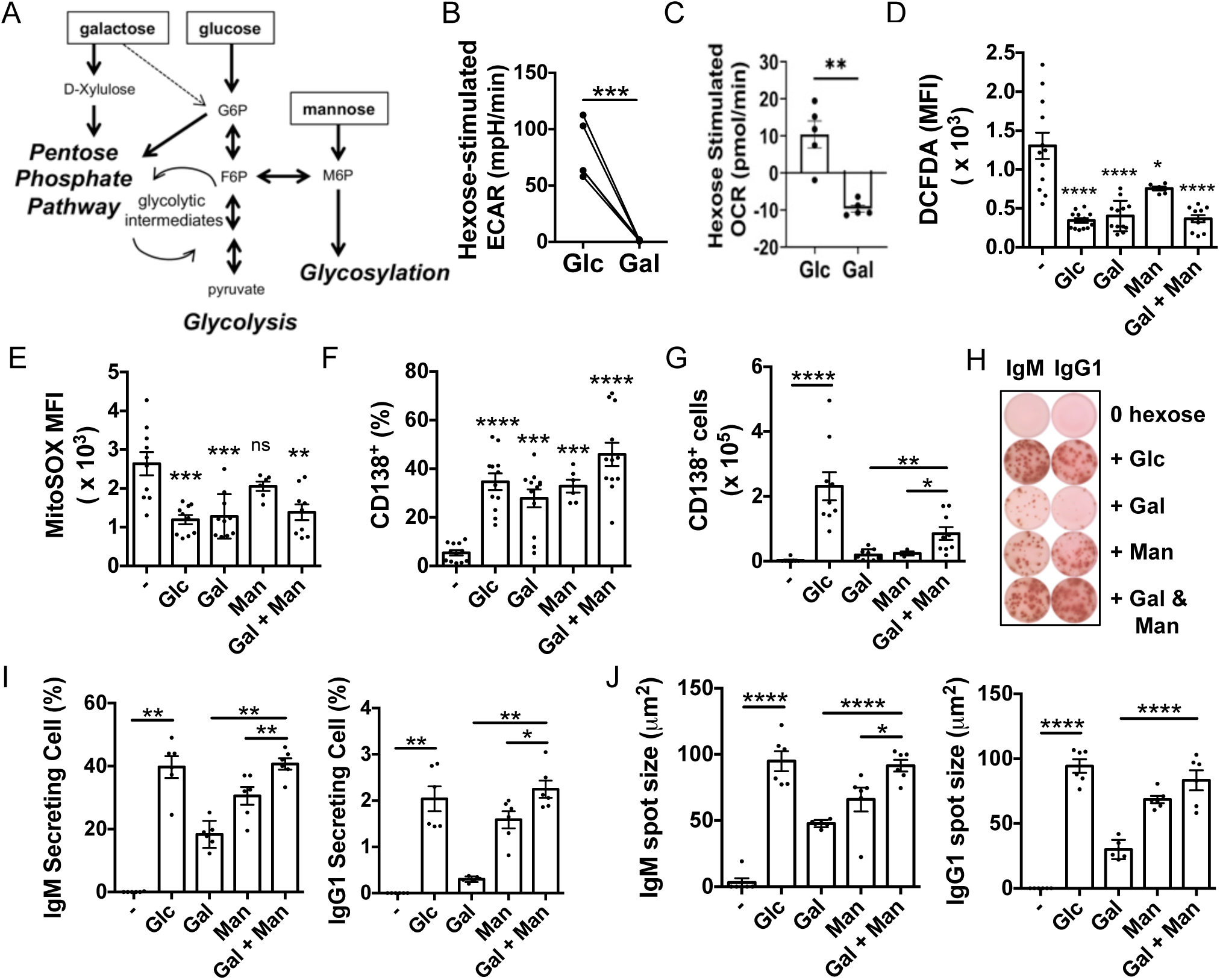
Collaborative hexose usage supports plasma cell and efficient antibody production. (A) Simplified schematic of hexose utilization and monosaccharide metabolism. (B) ECAR and (C) OCR measurements of LPS-activated wild-type B cells cultured (2 d) in BAFF, IL-4, IL-5. Metabolic fluxes of equal numbers of B lymphoblasts were analyzed using either glucose or galactose, as indicated, in ‘glycolytic stress testing’ programmed into the Seahorse XFe. Shown are the hexose-stimulated proton secretion (B), net of basal ECAR in hexose-free base medium, and mitochondrial respiration (C). Data represent four independent experiments with paired t-test comparisons. Representative raw data from a single experiment are shown in Supplemental Fig 3, panels C, D. (D) MFI of total cellular ROS (H_2_DCFDA) and (E) mitochondrial ROS (MitoSOX) of wild-type B lymphoblasts after activation with LPS and culture (2 d) with BAFF, IL-4, and IL-5 in glucose-free medium supplemented with glucose (Glc), galactose (Gal), Mannose (Man), or both Gal and Man. No hexose, (-). (F) Frequency of CD138^+^ cells in the live gate determined by flow cytometry after 5 days of culture as above. Results of statistical analyses in (D-F) represent comparison of each condition to the unsupplemented group by the Student’s t-test, with * - **** defined as in Fig 7 (G) Total number of CD138^+^ cells after provision of hexoses, calculated from the viable cell numbers and frequencies of CD138^+^ cells measured in (F). (H) Representative ELISpot wells of IgM and IgG1-secreting cells 5 days after cells were cultured in different hexose supplemented media as above and seeding equal numbers of viable cells (100 cells per well). (I) Quantitation of (H), depicting percentage of IgM and IgG1 secreting cells detected among total cells seeded in 5-day hexose cultures. (J) IgM and IgG1 antibody production per cell was analyzed by measurement of average spot sizes in each well. Data aggregate two independent replicate experiments, each with three independent B cell preparations cultured under the indicated conditions and analyzed by ELISpot assays. (G-J) Mann-Whitney U test determined statistical significance and indicated by * p < 0.05, ** p < 0.01, *** p < 0.001, **** p < 0.0001.

Proper glycosylation of antibodies is critical for plasma cell secretion and survival (56). Prior work indicated that most glucose use by plasma cells in media devoid of mannose was dedicated to glycosylation (9). However, mannose circulates in mammalian plasma at concentrations suited to uptake, likely in part through specialized transporters (57, 58). This hexose can be converted to mannose-6-phosphate (M6P), which then can either be transferred into glycolysis via fructose-6-phosphate (F6P), or used for glycosylation (59). I*n vivo* as well as *in vitro* studies have shown that N-linked glycosylation draws more on mannose than glucose as a precursor (7, 58). Accordingly, we tested if mannose could mitigate the impact of glucose-free conditions on redox balance, PC differentiation, and antibody production *in vitro*. Mannose supplementation did not dampen intracellular ROS or mtROS levels to the same degree as glucose or galactose, suggesting that contribution of mannose to PPP activity is less than the contributions of glucose or the Leloir pathway (Fig. 8D, E).

To test if glucose is uniquely essential for ASC differentiation and Ab production, we analyzed B cell differentiation into CD138^+^ cells by flow cytometry after *in vitro* culture with different hexose supplementation. Along with measuring rates of cell cycling in these conditions (Supplemental Fig. 4A, B) as well as frequencies and numbers of ASC, IgM and IgG1 in the culture supernatants, we estimated rates at which the differentiated cells secreted Ab by determination of spot sizes in ELISpot assays (Fig. 8F-J; Supplemental Fig. 4C, D). The frequency of CD138^+^ cells was restored regardless of which hexose was added to glucose-free media (Fig. 8F), but the total number of CD138^+^ cells in glucose-supplemented cultures was 10-fold higher than if galactose or mannose alone was provided (Fig. 8G). Of note, supplementation with both galactose and mannose produced more CD138^+^ cells compared to either hexose alone, suggesting that the requirement for glucose in plasma cell production could in part be bypassed when fueled with one or several hexoses that support PPP activity and glycosylation.

Furthermore, as determined by spot prevalence and sizes in ELISpot analyses at the end of cell cultures combined supplementation with galactose and mannose led to equally frequent and functional IgM- and IgG1-secreting cells compared to glucose-replete cultures (Fig. 8H-J). In contrast, neither galactose nor mannose could, on its own, restore net proliferation or Ab production in glucose-free conditions. Collectively, these data provide evidence of a preferential capacity of glucose to promote survival during differentiation of activated B into CD138^+^ antibody secreting cells. Surprisingly, however, this unique hexose requirement could be substantially albeit incompletely circumvented by a combination of alternate hexoses to support redox homeostasis by fueling the PPP (galactose) with a physiologically relevant means of supporting glycosylation (mannose). Thus, yields of CD138^+^ ASCs increased ∼1/3 as much as with glucose and, among the surviving population, these cells achieved robust Ab production without provision of glucose.

## DISCUSSION

Increasing evidence suggests that successful therapeutic targets may emerge from the elucidation of how specific metabolic pathways in lymphocytes alter their fate potential or functions (30, 60–62). Relative to T cells – or for that matter macrophages and DC - there are fewer stage-specific insights into how specific nutrients are used by B lymphocytes as they proceed toward constitutive Ab secretion as plasma cells (1, 2). Glucose metabolism generally constitutes one of the central networks for energetics, redox homeostasis, and provision of biosynthetic building blocks involving carbon. Based on experiments largely performed in vitro to complement in vivo observations, glucose is thought to play important roles in support of B cell activation and differentiation (11–13, 16), perhaps via provision of pyruvate for mitochondrial oxidation and metabolism (4, 31). Nonetheless, these processes were normal in experiments using glucose-free medium supplemented with regular FBS (35), and this concept has been challenged further based on observation of relatively modest completion of glycolysis and extracellular acidification by GC B cells assayed ex vivo (8).

We have shown herein that after normal differentiation and establishment of a pre-immune B cell repertoire the glucose transporter GLUT1 promotes initial GC B cell population numbers, affinity maturation, and PC outputs. ASC generation was reduced by a GC-specific approach to deleting *Slc2a1* coding sequences even after initiation of the GC B cell gene expression program and secondary follicle. Once taken up, glucose has multiple fates in the B lymphocyte lineage including the generation of nucleotides via the pentose phosphate pathway (PPP), lipid synthesis, fueling the TCA cycle, and the glycosylation of antibodies (3, 9, 35, 63). The findings presented here indicate that a polyclonal repertoire of purified GC B cells increased their rates of glucose uptake and oxidation (i.e., conversion to pyruvate and then acetyl-CoA) relative to naïve B counterparts, and that this increase was similar in magnitude to that measured with in vitro B lymphoblasts. However, in naïve, GCB, and blasting B cells the flux of glucose into the PPP was substantially greater than that of completing glycolysis, and whereas PPP activity increased yet more with development into PCs, this increase was prevented by GLUT1 depletion. The increased PPP function appeared crucial for quantitative outputs of PC after activation due to a capacity to promote redox homeostasis, but restoration of normal mitoSox and DCFDA signals did not suffice for normal outputs. Instead, provision of mannose together with PPP rescue was required for substantial output of ASC, after which Ab secretion could be normal in the absence of exogenous glucose. This surprising finding reveals ASC flexibility in adapting to nutrient limitation and sheds new light on mechanistic requirements for Ab responses.

Glucose is a major carbon source for activated and proliferating lymphocytes, but is it uniquely special for B lineage cells and if so why? Glucose is considered to be the most efficient of the simple sugars for fueling glycolysis; galactose preferentially is shuttled into the Leloir pathway, while mannose is directed to the glycosylation of proteins (53). Lymphocytes take up glucose down a concentration gradient through a family of GLUT transporters, which increase expression upon activation (3, 12, 37, 63, 64). *Cd19*-Cre-driven loss of GLUT1 during B cell development led to a substantial decrease in B cell numbers in the pre-immune state, and lowered an antibody response to immunization to the same extent as the cell population (12). This finding has been extended here by using cell type-specific and temporally-induced *Slc2a1* gene inactivation in mature B cells either prior to or after immunization. Along with results from in vitro analyses in which glucose was completely eliminated from cell cultures, the findings indicate that glucose influx rates affect affinity maturation and are crucial for outputs of Ab-secreting cells after B cell activation, including GC-derived PC.

Activated B cells subjected to depletion of GLUT1 - a plasma membrane carrier that mediates passive transport of glucose as well as other simple carbohydrates – after normal differentiation exhibited reduced rates of proliferation along with decreased ECAR. These findings suggest there is a threshold for glucose influx, below which optimal metabolic function is not achieved. Substantial proliferation precedes formation of the GC, whose maintenance depends on the balance among proliferation, death, and differentiation to MBC or PC [(65, 66); reviewed in (67)]. Conditionally GLUT1-deficient GC B cells were substantially reduced in an anti-SRBC response or when placed in competition with wild-type cells for immunizations with NP-ovalbumin (Fig. 4C and Fig. 3C, respectively), whereas in the absence of competitor B cells the defect was more modest in the early GC elicited by the same haptenated carrier (Fig. 3F) or an anti-protein recall response (Supplemental Fig. 2B). Precedents for such differential effects have been reported with altered mTORC1 activity or selective manipulation of dominant versus sub-dominant epitopes (68–71). These differences in degrees of impact (total GC B populations after SRBC versus NP-ova; NP-specific versus total GC B cells are likely due to a combination of several factors. First, epitope valency alters the robustness of GC development, maintenance, and outputs (72). Moreover, the dynamics of competition for limiting T cell help in GC populations (73–75) means that a reduction in proliferative rate can be mitigated by lower competition and thus more survival. The findings may also arise if there is increased uptake by other hexose transporters that could compensate for the loss of *Slc2a1* and as a consequence of the variable degree of counter-selection favoring the small minority of B cells without recombination of both fl alleles (Fig. 3G; Fig. 4E). In any event, the data with *S1pr2*-Cre-driven deletion, taken together with in vitro experiments, indicate that the effectiveness of GC outputs into the ASC population is reduced when glucose influx is constrained.

*Slc2a1* inactivation caused a strong defect in plasma cell differentiation and antibody production both *in vitro* and in response to immunization, in the latter case even when conditional loss-of-function was initiated with the GC-selective *S1pr2*-CreER^T2^ gene. These findings suggest that demands for glucose of plasma cells and/or their differentiation exceed those of GCB, consistent with the heightened 2-NBDG labeling of ASC compared to other B cells subsets and prior work of others (9, 10). An intriguing aspect of the data is that GLUT1 depletion reduced the ability to select or produce high-affinity anti-NP Ab. Inasmuch as higher BCR affinity biases or selects for progression to PC fate (76–79), it is possible - but not tested here – that the observed reductions in PC output after *Slc2a1* inactivation is due at least in part to less efficient generation of higher-affinity BCRs.

Our findings establish that adequacy of glucose influx fosters enhanced metabolic rates after B cell activation, so that both the rates of extracellular acidification and respiration were less robust with *Slc2a1* Δ*/*Δ B cells. Another function is to supply the PPP, whose measured activity was higher than glucose oxidation, which depends on glycolytic flux to pyruvate. Prior in vitro work has used isotopomer metabolite analyses elegantly to show that in vitro B lymphoblasts diverted most of their glucose to the PPP (3, 35). The tracing studies further indicated that a substantial fraction of glucose was used for nucleotide synthesis in the first 24-48 h after B cell activation with anti-CD40/IL-4 or anti-IgM (3, 35). The work here extends these insights to fresh ex vivo GC B cells, and provides evidence that the pentose phosphate ‘shunt’ pathway increases the efficiency of PC production in vitro.

Our findings indicate that an adequate influx of hexose through GLUT1 is important for maintenance of ROS homeostasis, presumably via the PPP. However, apparent normalization of ROS under glucose-free conditions (i.e., via provision of galactose) did not suffice for B cells to survive in differentiating, so that glucose or some other hexose is needed for additional pathway(s). Although the PPP may also be important because it can contribute to the cell-intrinsic generation of nucleoside precursors to support growth, analyses of transformed B cells support a crucial role for glucose in nucleotide synthesis. Of note, GLUT1 was critical for the synthesis of multiple metabolites involved in the PPP and pyrimidine/purine metabolism in BCR-Abl^+^ B-cell acute lymphoblastic leukemia cells (B-ALL) (80). Moreover, *de novo* nucleotide synthesis is crucial for the proliferation and cell cycle progression of activated T lymphocytes (81). In light of the published isotopomer data with activated B cells (35), it is likely that proliferating B cells also need cell-intrinsic genesis of nucleotide precursors. Consistent with this, protein phosphatase 2A-mediated maintenance of an appropriate balance between glycolysis and PPP and sufficient flux through the PPP was reported to promote B cell and B-ALL survival (49). In principle, ROS scavengers such as N-acetylcysteine or cell-permeable glutathione might aid survival in the hexose-free medium. Unfortunately, these agents grossly perturbed outputs (cell numbers; % CD138^+^) even in normal glucose-replete medium (C. Paik, unpublished observations). Thus, while a causal connection of the capacity of galactose to normalize cytosolic and mitochondrial ROS and its capacity to collaborate with mannose in partial rescue of PC production seems most likely because of the limited alternatives, that causality is not fully established by the data.

In addition to usage by the PPP, however, additional requirements for sufficient hexose can be inferred from our data. One is the need to support protein glycosylation, N-linked glycosylation of Ab is essential for avoiding cell death due to ER stress (82, 83). In tracer experiments using medium devoid of mannose, a large fraction of glucose-derived ^14^C was found in N-linked glycans on Ab (9). Strikingly, however, the in vitro-generated PC that arose in the absence of glucose while cultured in the presence of mannose along with galactose generated spot sizes indistinguishable from the spots generated by secretion from glucose-replete control cultures. Glycosylation can affect cell viability and efficiencies of Ab secretion (82–84), so the results suggest that metabolic flexibility in ASCs allows mannose to mitigate the effect that glucose deprivation has on generation of glycanated Ab. Conversely, since direct in vivo and in vitro data have indicated that mannose is the main physiological source of sugars for glycosylation (57, 58), the finding raises the possibility that in vivo mannose may be the main hexose used for glycosylation of Ab. Finally, the data suggest that mannose supply protected from death. For net proliferation and survival to reach the CD138^+^ stage, however, the output of ASCs was higher in glucose-supplemented medium than with mannose and galactose in glucose-free conditions. Accordingly, glucose appeared to make at least one other, as yet unidentified, biochemical contribution to the capacity to achieve normal Ab outputs. W infer that insufficient glucose influx cannot be replaced by circulating galactose and mannose in vivo because of a pathway fed by glucose but not these epimers. However, further work will be required to elucidate this function and test the model that this usage is an additional mechanism contributing to the defects of GLUT1-deficient B lineage cells.

## ACKNOWLEDGEMENTS

We thank J. Cyster for generously shipping *S1pr2*-CreERT2 breeding stock, M. Reth for suggesting the galactose method of providing substrate for the PPP, A. D. Posey Jr. for support to S. K. B. during manuscript preparation and revision, and Vanderbilt institutional cores (High-Throughput Screening; Flow Cytometry Shared Resource; Small Molecule NMR; Cell & Developmental Biology) for equipment, expertise, and assistance.

## FOOTNOTES

Experimental work was supported by NIH grant R01 AI149722 (M. R.B), with metabolome analyses supported by R01 AI153167 and DK105550 (J. C. R.). Additional support for S. K. B. was provided by NIH grants R25 GM062459, T32 CA009592-29, and a Provost Graduate Fellowship (Vanderbilt University), followed by departmental funds (P. M. & I.). F31 HL152529 provided training support to D. L. G., and NIH Shared Instrumentation Grant 1S10OD018015 as well as scholarships via the Cancer Center Support Grant (CA068485) and Diabetes Research Center (DK0205930) helped defray costs of Vanderbilt Cores.

## Abbreviations Used

Ab: antibody
ASC: Ab-secreting cell
GC: germinal center
PC: plasma cell
LLPC: long-lived PC
2DG: 2-deoxyglucose
2-NBDG: 2-(N-Nitrobenz-2-oxa-1,3-diazol-4-yl)amino)-2-deoxyglucose
FCCP: Trifluoromethoxy carbonylcyanide phenylhydrazone
PPP: Pentose Phosphate Pathway
ROS: reactive oxygen species
mTORC1: mechanistic target of rapamycin complex 1
mtROS: mitochondrial ROS
ECAR: extracellular acidification rate
OCR: oxygen consumption rate
NP: 4-hydroxy-3-nitrophenylacetyl (nitrophenyl)
NP-ova: NP conjugated to ovalbumin
4-OHT: 4-hydroxytamoxifen
CTV: CellTrace Violet
Glc: glucose
Gal: galactose
Man: mannose

## SUPPLEMENTAL DATA

**Supplemental Figure 1.**
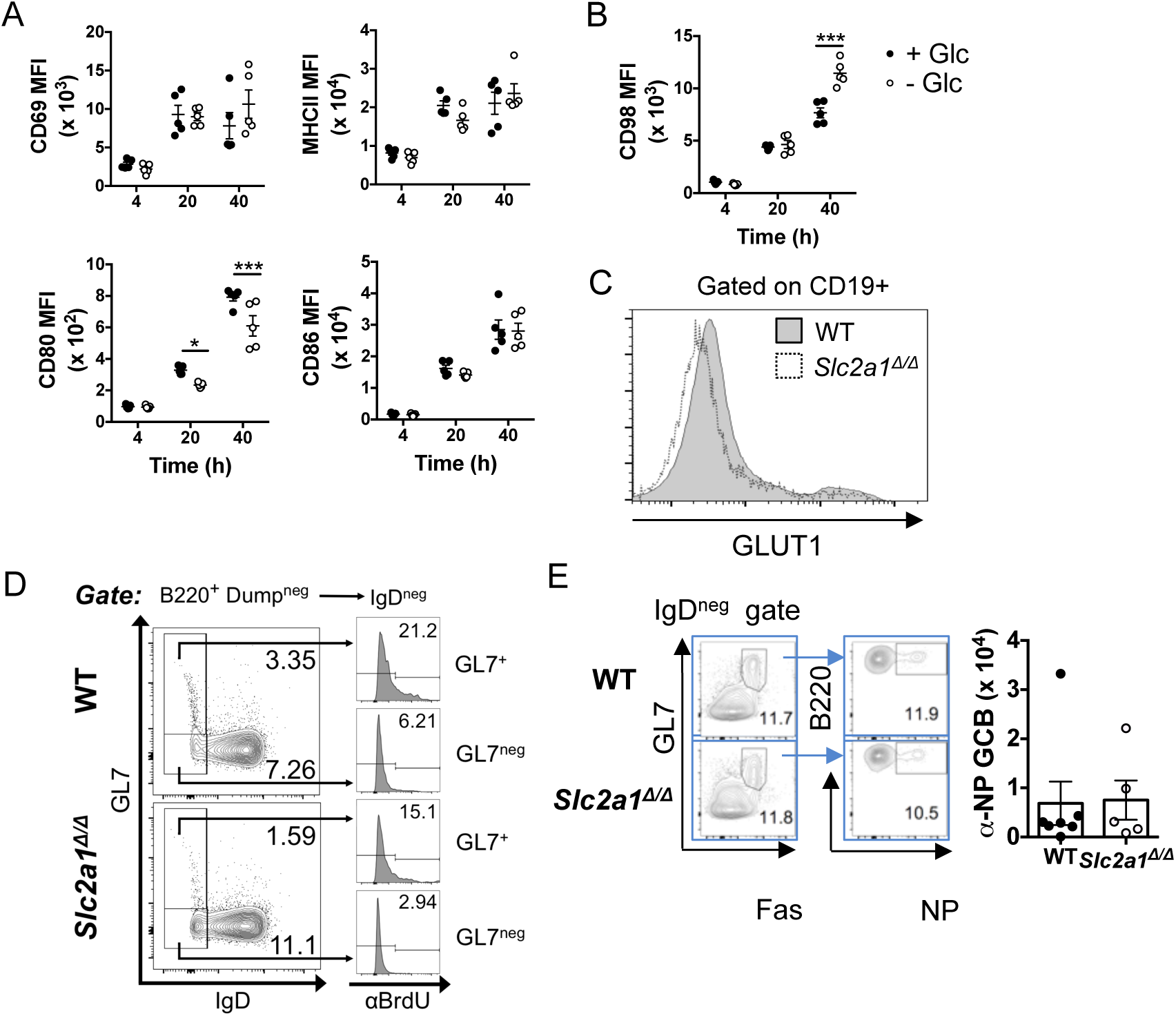
Early activation markers induced in absence of glucose. (A) CD69, MHC-II, CD80, and CD86 surface expression on wild-type B cells after activation by LPS and culture (4, 20, and 40 hr, as indicated) in BAFF, IL-4, and IL-5, in the absence or presence of glucose, as indicated. (B) CD98 surface expression on B cells 4, 20, and 40h after LPS, BAFF, IL-4, and IL-5 stimulation. Shown are the aggregate results of two independent experiments with five total replicates. * p < 0.05, *** p < 0.001 (Multiple comparisons t-tests). (C) Surface GLUT1 expression is diminished on *Slc2a1* Δ*/*Δ (cKO) B cells. Representative flow plot of surface GLUT1 (Alomone Lab) on CD19^+^ B cells were from tamoxifen-treated WT (*hCD20*-CreER^T2^) and Glut1 cKO^B^ (*Slc2a1^f/f^*; *hCD20*-CreER^T2^) mice. (D) Frequencies of BrdU^+^ cells in IgD^−^ GL7^+^ CD38^neg^ B cells. Tamoxifen-treated WT (*hCD20*-CreER^T2^) and Glut1 cKO^B^ (*Slc2a1^f/f^*; *hCD20*-CreER^T2^) mice were immunized with SRBC and harvested 7 days later. Mice were injected twice with BrdU (100 mg/kg i.v.), 1 d and 4 h before harvest.

**Supplemental Figure 2.**
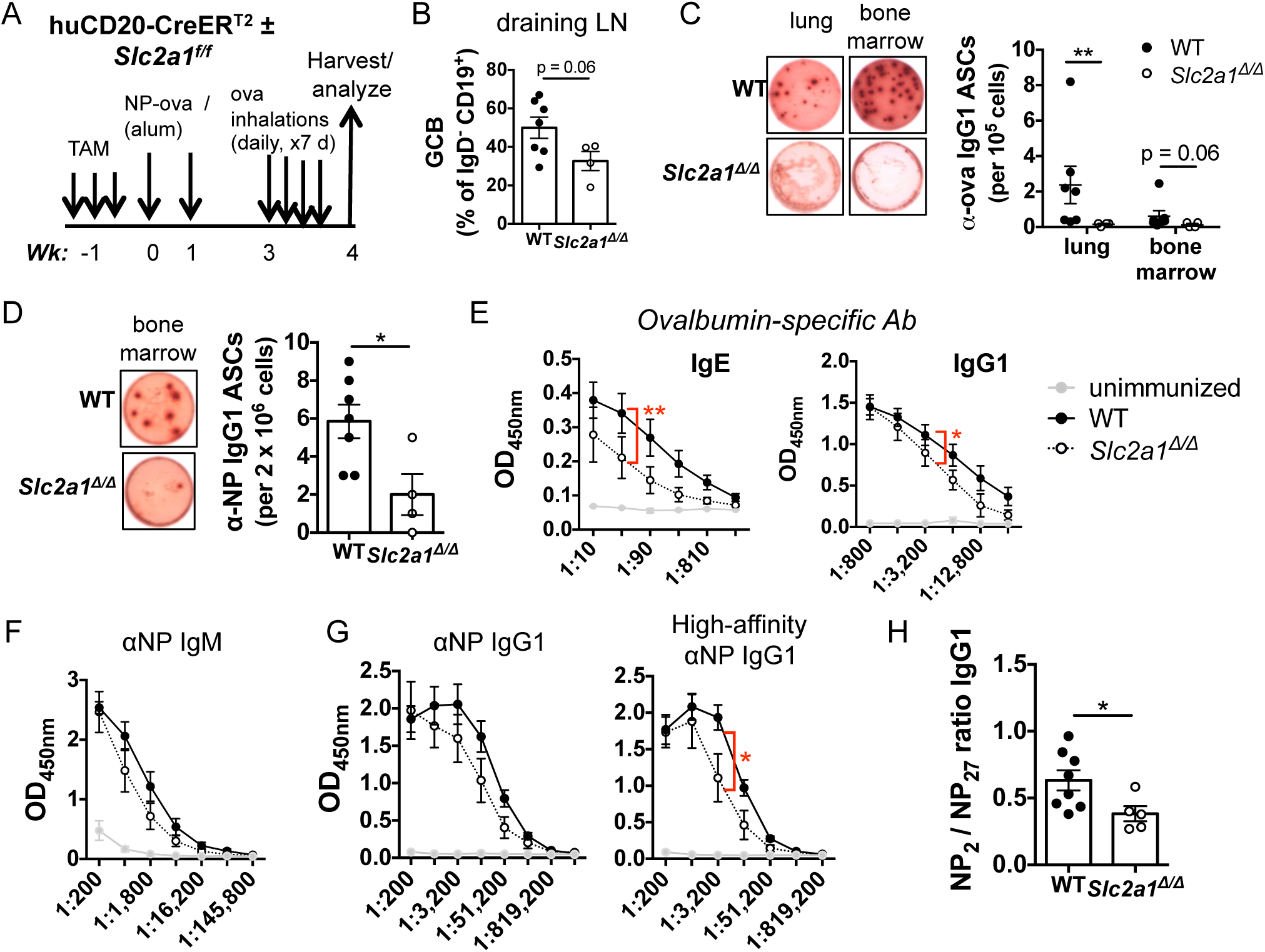
GLUT1 in B cells supports the generation of ASC and high affinity antibodies in a lung inflammation model. (A) Schematic of immunization strategy to test B cell loss of *Slc2a1* in a lung inflammation model. Mice were primed with two I.P. injections of NP-ova, followed by sequential ovalbumin inhalations received daily for seven days starting 3 wk after initial immunization and continuing until harvest. (B) Frequency of GCB in mediastinal lymph nodes of wildtype and *Slc2a1^Δ/Δ^* mice at the time of harvest, as determined by flow cytometry. (C) *Left panel*, representative ELISpot wells of ovalbumin-specific IgG1-secreting cells in the lung (among 10^6^ cells in lung dispersion) and bone marrow (2 x 10^6^ bone marrow cells plated/well) of mice with wild-type or *Slc2a1^Δ/Δ^* B cells. *Right panel*, numbers of ovalbumin-specific IgG1 antibody-secreting cells in the lung or bone marrow per 10^5^ total cells. (D) As in (C) except ELISpot captured NP-specific ASC. *Left panel*, representative ELISpot wells visualizing IgG1-secreting bone marrow cells from immunized mice with wild-type or *Slc2a1^Δ/Δ^* B cells. *Right panel*, numbers of NP-specific IgG1 ASC in the bone marrow per 10^6^ cells. (E) Ovalbumin-specific IgE (left panel) and IgG1 (right panel) in unimmunized, wild-type, and *Slc2a1^Δ/Δ^* mice at the terminal time point depicted as OD_450nm_ values after three or two-fold serial dilutions of the sera for ELISA, respectively. (F) NP-specific IgM in the sera of mice at week 3, prior to inhaled challenges with ovalbumin. As for (E), except serial dilutions were 4-fold. (G) All-affinity (left panel) and high affinity (right panel) NP-specific IgG1 in the sera of mice at week 3, prior to inhaled challenges with ovalbumin. (H) Shown are the ratios of OD values of NP_2_-bound IgG1 antibodies (high affinity) measured in ELISA to those of NP_27_-bound (all affinity) IgG1; ratios were calculated at a dilution (1:12,800) with values for both WT and GLUT1-deficient B cells in the linear range. Data are representative of two independent experiments with n = 7 wild-type and n = 4 *Slc2a1^Δ/Δ^* mice. * p < 0.05, ** p < 0.01. Mann Whitney U (B-D, H), Two-way ANOVA (E, G).

**Supplemental Figure 3.**
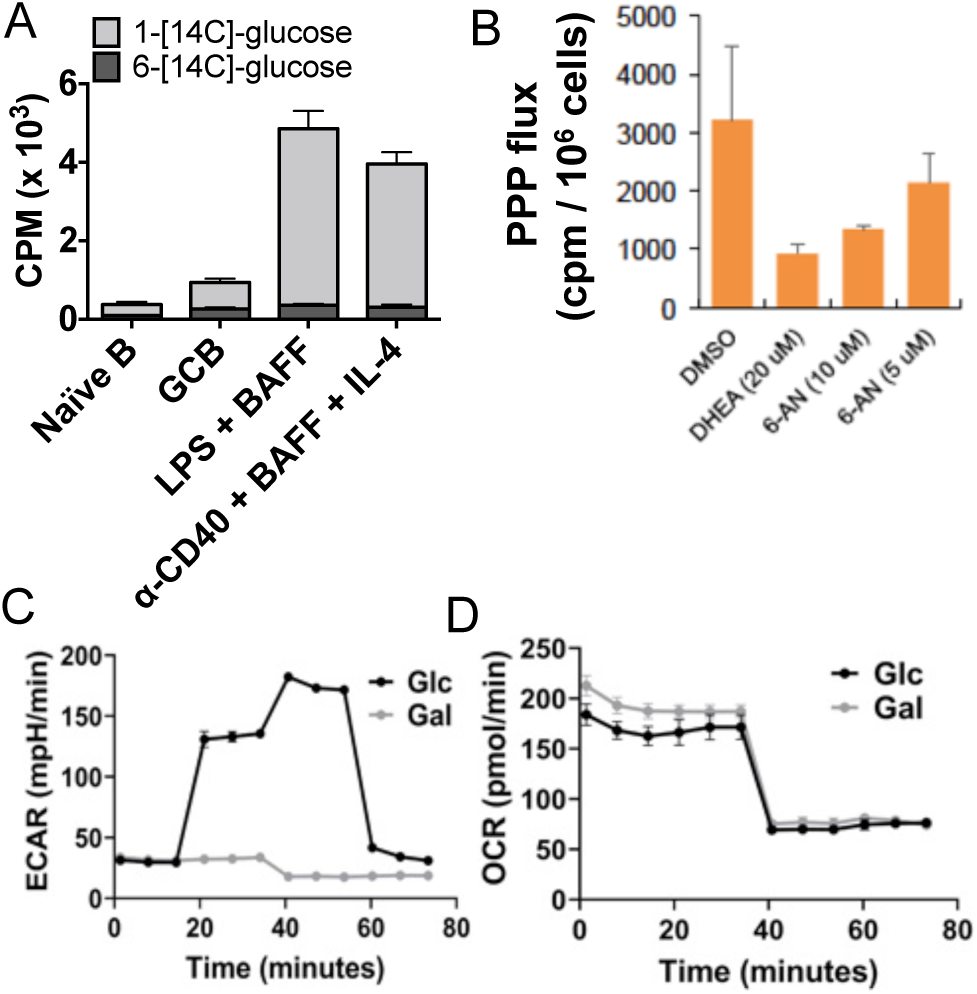
Glucose is fated for PPP and oxidation in the B lineage and combined metabolism of hexoses supports antibody production. **(A)** PPP activity and glucose oxidation determined by 1-[^14^C]-glucose and 6-[^14^C]-glucose conversion to ^14^CO_2_ in the indicated flow sorted B cell populations after SRBC immunization or *in vitro* activated B lymphoblasts (LPS and BAFF or anti-CD40, BAFF, and IL-4, as indicated). (B) PPP activity, measured using 1-[^14^C]-glucose, in wild-type B cell cultures two days after activation (anti-CD40) and culture as in (A), and treatment with oxidative PPP inhibitors, DHEA and 6-AN as indicated. Galactose generates neither acidification (C) nor respiration (D) by CD40-activated B lymphoblasts. Shown are the full tracings of measurements using the Seahorse xFe analyzer in one experiment representative of four biologically independent replications. As in Fig. 8C, D, purified splenic B cells were activated with LPS, cultured 2 d with BAFF, IL-4, and IL-5, rinsed, counted, and aliquoted for metabolic flux assays in base medium for glycolytic testing, with one set of triplicate samples receiving glucose and a parallel set galactose after measurement of the basal metabolism.

**Supplemental Figure 4.**
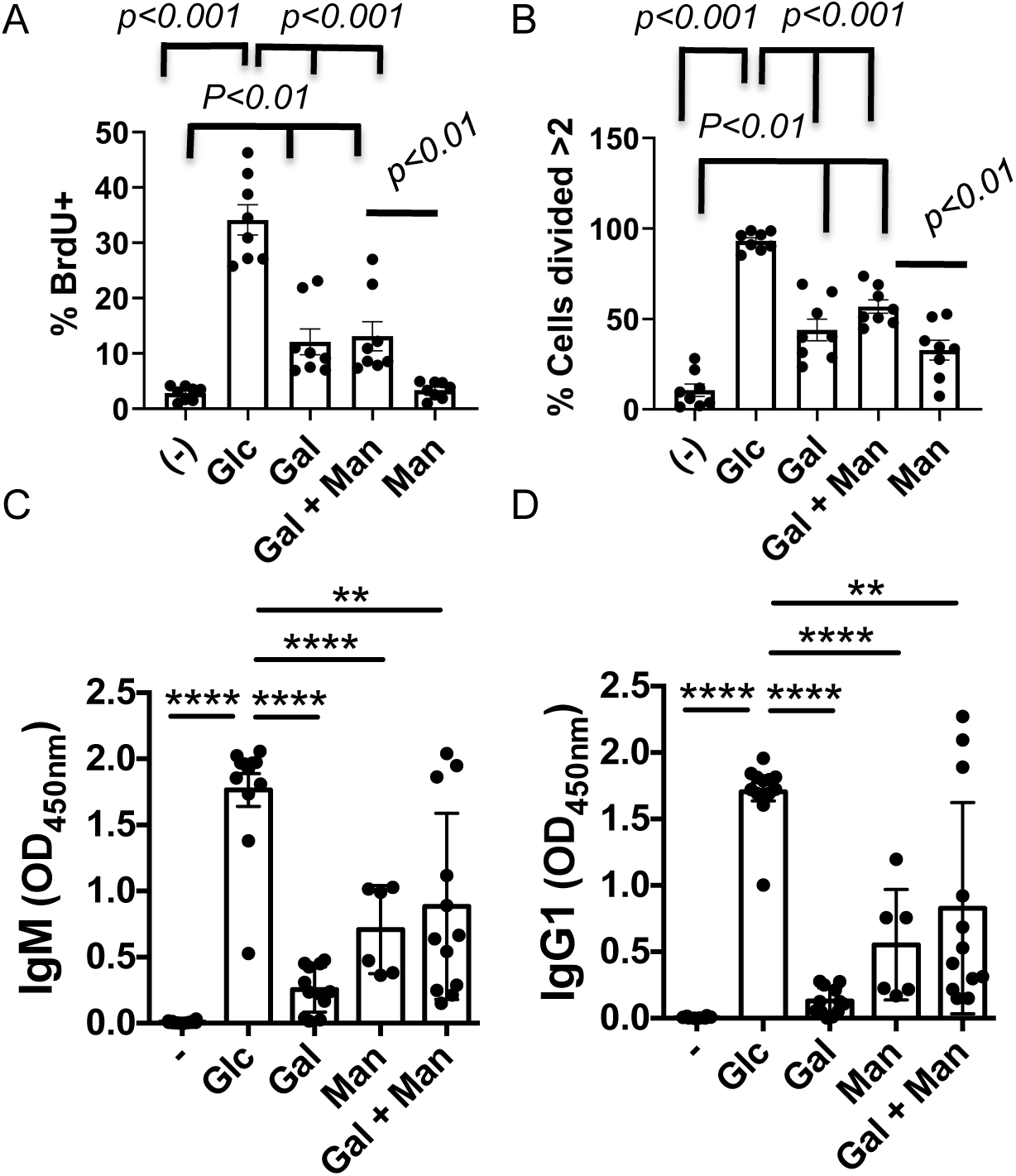
Glucose is fated for PPP and oxidation in the B lineage and combined metabolism of hexoses supports antibody production. Cell cycling (A) and division (B) of activated B cells partially supported by galactose in the absence of exogenous glucose. Purified splenic B cells were activated with LPS, aliquoted, and culture with BAFF, IL-4, and IL-5 in glucose-free medium supplemented with dialyzed FBS and the indicated hexose(s). The resulting populations were analyzed by BrdU pulse-labeling and flow cytometry after two (A) days, or (B) by measurement of the division counts based on decreases in CTV fluorescence after four days. (C, D) Ab concentrations tracked with PC population sizes when galactose and mannose substituted for glucose. IgM (C) and IgG (D) Ab accumulated in the culture supernatants from the experiments in Figure 8, F-J. Purified splenic B cells were activated with LPS, aliquoted, and cultured 5 d with BAFF, IL-4, and IL-5 in glucose-free medium supplemented with dialyzed FBS and the indicated hexose(s). IgM and IgG1 in the supernatant after 5 days of culture were measured by ELISA.

## Notes

### Competing Interest Statement

The authors have declared no competing interest.

## REFERENCES

1. Ripperger, T. J. and D. Bhattacharya. 2021. Transcriptional and metabolic control of memory B cells and plasma cells. Annu Rev Immunol. 39: 345–368.

2. Boothby, M.R., S. K. Brookens, A. L. Raybuck, and S. H. Cho. 2022. Supplying the trip to antibody production—nutrients, signaling, and the programming of cellular metabolism in the mature B lineage. Cell Mol Immunol 19: 352–369.

3. Doughty, C.A., B. F. Bleiman, D. J. Wagner, F. J. Dufort, J. M. Mataraza, M. F. Roberts, and T. C. Chiles. 2006. Antigen receptor-mediated changes in glucose metabolism in B lymphocytes: role of phosphatidylinositol 3-kinase signaling in the glycolytic control of growth. Blood 107: 4458–4465.

4. Cho, S. H., A. K. Ahn, P. Bhargava, C.-H. Lee, C. M. Eischen, O. McGuinness, and M. Boothby. 2011. Glycolytic rate and lymphomagenesis depend on PARP14, an ADP ribosyltransferase of the B aggressive lymphoma (BAL) family. Proc Natl Acad Sci USA 108: 15972–15977.

5. Cho, S. H., A. L. Raybuck, K. Stengel, M. Wei, T. C. Beck, E. Volanaki, J. W. Thomas, S. Hiebert, V. H. Haase, and M. R. Boothby. 2016. Germinal centre hypoxia and regulation of antibody qualities by a hypoxia response system. Nature 537: 234–237.

6. Jellusova, J., M. H. Cato, J. R. Apgar, P. Ramezani-Rad, C. R. Leung, C. Chen, A. D. Richardson, E. M. Conner, R. J. Benschop, J. R. Woodgett, and R. C. Rickert. 2017. Gsk3 is a metabolic checkpoint regulator in B cells. Nat Immunol 18: 303–312.

7. Vijay, R., J. J. Guthmiller, A. J. Sturtz, F. A. Surette, K. J. Rogers, R. R. Sompallae, F. Li, R. L. Pope, J. Chan, F. L. Rivera, D. Andrew, L. Webb, W. J. Maury, H. H. Xue, C. R. Engwerda, J. S. McCarthy, M. J. Boyle, and N. S. Butler. 2020. Infection-induced plasmablasts are a nutrient sink that impairs humoral immunity to malaria. Nat Immunol 21: 790–801.

8. Weisel, F. J., S. J. Mullett, R. A. Elsner, A. V. Menk, N. Trivedi, W. Luo, D. Wikenheiser, W. F. Hawse, M. Chikina, S. Smita, L. J. Conter, S. M. Joachim, S. G. Wendell, M. J. Jurczak, T. H. Winkler, G. M. Delgoffe, and M. J. Shlomchik. 2020. Germinal center B cells selectively oxidize fatty acids for energy while conducting minimal glycolysis. Nat Immunol 21: 331–342.

9. Lam, W. Y., A. M. Becker, K. M. Kennerly, R. Wong, J. D. Curtis, E. M. Llufrio, K. S. McCommis, J. Fahrmann, H. A. Pizzato, R. M. Nunley, J. Lee, M. J. Wolfgang, G. J. Patti, B. N Finck, E. L. Pearce, and D. Bhattacharya. 2016. Mitochondrial pyruvate import promotes long-term survival of antibody-secreting plasma cells Immunity 19: 60–73.

10. Lam, W. Y., A. Jash, C. H. Yao, J. D’Souza, R. Wong, R. M. Nunley, G. P. Meares, G. J. Patti, D., and D. Bhattacharya. 2018. Metabolic and transcriptional modules independently diversify plasma cell lifespan and function. Cell Rep 24: 2479–2492.

11. Kojima, H., A. Kobayashi, D. Sakurai, Y. Kanno, H. Hase, R. Takahashi, Y. Totsuka, G. L. Semenza, M. V. Sitkovsky, and T. Kobata. 2009. Differentiation stage-specific requirement in hypoxia-inducilbe factor-1**α**-regulated glycolytic pathway during murine B cell development in bone marrow. J Immunol 184; 154–163.

12. Caro-Maldonado, A., R. Wang, A. G. Nichols, M. Kuraoka, S. Milasta, L. D. Sun, A. L. Gavin, E. D. Abel, G. Kelsoe, D. R. Green, and J. C. Rathmell. 2014. Metabolic reprogramming is required for the antibody production that is suppressed in anergic but exaggerated in chronically BAFF-exposed B cells. J Immunol 192: 3626–3636.

13. Urbanczyk, S., M. Stein, W. Schuh, H. M. Jäck, D. Mougiakakos, and D. Mielenz. 2018. Regulation of energy metabolism during B lymphocyte development. Int J Mol Sci 19: 1–16.

14. Bertolotti, M., S. H. Yim, J. M. Garcia-Manteiga, S. Masciarelli, Y. J. Kim, M. H. Kang, Y. Iuchi, J. Fujii, R. Vené, A. Rubartelli, S. G. Rhee, and R. Sitia. 2010. B- to plasma-cell terminal differentiation entails oxidative stress and profound reshaping of the antioxidant responses. Antioxid Redox Signal 13: 1133–44.

15. Muri, J., H. Thut, S. Heer, C. C. Krueger, G. W. Bornkamm, M. F. Bachmann, and M. Kopf. 2019. The thioredoxin-1 and glutathione/glutaredoxin-1 systems redundantly fuel murine B-cell development and responses. Eur J Immunol 49: 709–723.

16. Donnelly, R. P., and D. K. Finlay. 2015. Glucose, glycolysis and lymphocyte responses. Mol Immunol 2015; 68:513–9.

17. Toretta, S., A. Scagliola, L. Ricci, F. Mainini, S. D. Marco, I. Cuccovillo, A. Kajaste-Rudnitski, D. Sumpton, K. M. Ryan, and S. Cardaci. 2020. D-mannose suppresses macrophage IL-1β production. Nat Commun 11: 6343.

18. Young, C. D., A. S. Lewis, M. C. Rudolph, M. D. Ruehle, M. R. Jackman, U. J. Yun, O. Ilkun, R. Pereira, E. D. Abel, and S. M. Anderson. 2022. Modulation of glucose transporter (GLUT1) expression levels alters mouse mammary tumor cell growth in vitro and in vivo. PLoS One 6: e23205.

19. Ahuja, A., J. Shupe, R. Dunn, M. Kashgarian, M. R. Kehry, and M. J. Shlomchik. 2007. Depletion of B cells in murine lupus: efficacy and resistance. J Immunol 179: 3351–61.

20. Raybuck, A. L., S. H. Cho, J. Li, M. Rogers, K. Lee, C. L. Williams, M. Shlomchik, J. W. Thomas, J. Chen, J. V. Williams, and M. R. Boothby. 2018. B cell-intrinsic mTORC1 promotes GC-defining transcription factor gene expression, somatic hypermutation, and memory B cell generation in humoral immunity. J. Immunol 200: 2627–2639.

21. Shinnakasu, R., T. Inoue, K. Kometani, S. Moriyama, Y. Adachi, M. Nakayama, Y. Takahashi, H. Fukuyama, T. Okada, and T. Kurosaki. 2016. Regulated selection of germinal-center cells into the memory B cell compartment. Nat Immunol 17: 861–869.

22. Brookens, S. K., S. H. Cho, P. J. Basso, and M. R. Boothby. 2020. AMPKα1 in B cells dampens primary antibody responses yet promotes mitochondrial homeostasis and persistence of B cell memory. J Immunol 205: 3011–3022.

23. Schmidt-Supprian, M. and K. Rajewsky. 2007. Vagaries of conditional gene targeting. Nat Immunol 8: 665–668.

24. Becher, B., A. Waisman, and L. Lu. 2018. Conditional gene-targeting in mice: problems and solutions. Immunity 48: 835–836.

25. Cho, S. H., A. L. Raybuck, J. Blagih, E. Kemboi, V. H. Haase, R. G. Jones, and M. R. Boothby. 2019. Hypoxia-inducible factors in CD4^+^ T cells promote metabolism, switch cytokine secretion, and T cell help in humoral immunity. Proc Natl Acad Sci USA 116: 8975–8984.

26. Lee, K., K. T. Nam, S. H. Cho, P. Gudapati, Y. Hwang, D. S. Park, R. Potter, J. Chen, E. Volanakis, and M. R. Boothby. 2012. Vital roles of mTOR complex 2 in Notch-driven thymocyte differentiation and leukemia. J Exp Med 209: 713–728.

27. Wang, R., C. P. Dillon, L. Z. Shi, S. Milasta, R. Carter, D. Finkelstein, L. L. McCormick, P. Fitzgerald, H. Chi, J. Munger, and D. R. Green. 2011. The transcription factor Myc controls metabolic reprogramming upon T lymphocyte activation. Immunity 35: 871–882.

28. Pescarmona, G. P., A. Bosia, P. Arese, M. L. Sartori, and D. Ghigo. 1982. A simplified method for the pentose phosphate pathway assay in red cells. Int J Biochem 14: 243–245.

29. Govindaraju, V., K. Young, and A. A. Maudsley. 2000. Proton NMR chemical shifts and coupling constants for brain metabolites. NMR Biomed 13:129–153.

30. Sugiura, A., G. Andrejeva, K. Voss, D. R. Heintzman, X. Xu, M. Z. Madden, X. Ye, K. L. Beier, N. U. Chowdhury, M. M. Wolf, A. C. Young, D. L. Greenwood, A. E. Sewell, S. K. Shahi, S. N. Freedman, A. M. Cameron, P. Foerch, T. Bourne, J. C. Garcia-Canaveras, J. Karijolich, D. C. Newcomb, A. K. Mangalam, J. D. Rabinowitz, and J. C. Rathmell. 2022. MTHFD2 is a metabolic checkpoint controlling effector and regulatory T cell fate and function. Immunity 55: 65–81.

31. Haniuda, K., S. Fukao, and D. Kitamura. 2020. Metabolic reprogramming induces germinal center B cell differentiation through Bcl6 locus remodeling. Cell Rep 33: 1–14.

32. Sinclair, L. V., C. Barthelemy, and D. A. Cantrell. 2020. Single cell glucose uptake assays: a cautionary tale. Immunometabolism 2: 1–13.

33. Hamilton, K. E., M. F. Bouwer, L. L. Louters, and B. D. Looyenga. 2021. Cellular binding and uptake of fluorescent glucose analogs 2-NBDG and 6-NBDG occurs independent of membrane glucose transporters. Biochimie 190: 1–11.

34. Reinfeld, B. I., M. Z. Madden, M. M. Wolf, A. Chytil, J. E. Bader, A. R. Patterson, A. Sugiura, A. S. Cohen, A. Ali, B. T. Do, A. Muir, C. A. Lewis, R. A. Hongo, K. L. Young, R. E. Brown, V. M. Todd, T. Huffstater, A. Abraham, R. T. O’Neil, M. H. Wilson, F. Xin, M. N. Tantawy, W. D. Merryman, R. W. Johnson, C. S. Williams, E. F. Mason, F. M. Mason, K. E. Beckermann, M. G. Vander Heiden, H. C. Manning, J. C. Rathmell, and W. K. Rathmell. 2021. Cell-programmed nutrient partitioning in the tumour microenvironment. Nature 593: 282–288.

35. Waters, L. R., F. M. Ahsan, D. M. Wolf, O. Shirihai, and M. A. Teitell. 2018. Initial B cell activation induces metabolic reprogramming and mitochondrial remodeling. iScience 5: 99–109.

36. Khalil, A. M., J. C. Cambier, and M. J. Shlomchik. 2012. B cell receptor signal transduction in the GC is short-circuited by high phosphatase activity. Science 336: 1178–1181.

37. Maratou, E., G. Dimitriadis, A. Kollias, E. Boutati, V. Lambadiari, P. Mitrou, and S. A. Raptis. 2007. Glucose transporter expression on the plasma membrane of resting and activated white blood cells. Eur J Clin Invest 37: 282–290.

38. Yazicioglu Y. F., E. Marin, C. Sandhu, S. Galiani, I. G. A. Raza, M. Ali, B. Kronsteiner, E. B. Compeer, M. Attar, S. J. Dunachie, M. L. Dustin, and A. J. Clarke. 2023. Dynamic mitochondrial transcription and translation in B cells control germinal center entry and lymphomagenesis. Nat Immunol. 24: 991–1006.

39 Silva, M., T. H. Nguyen, P. Philbrook, M. Chu, O. Sears, S. Hatfield, R. K. Abbott, G. Kelsoe, and M. V. Sitkovsky. 2017. Targeted elimination of immunodominant B cells drives the germinal center reaction toward subdominant epitopes. Cell Rep 21: 3672–3680.

40. Woodruff, M. C., Kim, E. H., Luo, W., and Pulendran, B. 2018. B cell competition for restricted T cell help suppresses rare epitope responses. Cell Rep 25: 321–327.

41. Brooks, J. F., C. Tan, J. L. Mueller, K. Hibiya, R. Hiwa, V. Vykunta, and J. Zikherman. 2021. Negative feedback by NUR77/Nr4a1 restrains B cell clonal dominance during early T-dependent immune responses. Cell Rep 36: 109645.

42. MacLennan, I. C., K. M. Toellner, A. F. Cunningham, K. Serre, D. M. Sze, E. Zuniga, M. C. Cook, and C. G. Vinuesa. 2003. Extrafollicular antibody responses. Immunol Rev 194: 8–18.

43. Hsu, M. C., K. M. Toellner, C. G. Vinuesa, and I. C. Maclennan. 2006. B cell clones that sustain long-term plasmablast growth in T-independent extrafollicular antibody responses. Proc Natl Acad Sci USA 103: 5905–5510.

44. Good-Jacobson, K. L., and M. J. Shlomchik. 2010. Plasticity and heterogeneity in the generation of memory B cells and long-lived plasma cells: the influence of germinal center interactions and dynamics. J Immunol 185: 3117–3125.

45. Viant, C., T. Wirthmiller, M. A. ElTanbouly, S. T. Chen, M. Cipolla, V. Ramos, T. Y. Oliveira, L. Stamatatos, and M. C. Nussenzweig. 2021. Germinal center-dependent and - independent memory B cells produced throughout the immune response. J Exp Med 218: e20202489.

46. Viant, C., G. H. J. Weymar, A. Escolano, S. Chen, H. Hartweger, M. Cipolla, A. Gazumyan, and M. C. Nussenzweig. 2020. Antibody affinity shapes the choice between memory and germinal center B cell fates. Cell 183: 1298–1311.

47. Green, J. A., and J. G. Cyster. 2012. S1PR2 links germinal center confinement and growth regulation. Immunol Rev 247: 36–51.

48. Dufort, F. J., M. R. Gumina, N. L. Ta, Y. Tao, S. A. Heyse, D. A. Scott, A. D. Richardson, T. N. Seyfried, and T. C. Chiles. 2014. Glucose-dependent de novo lipogenesis in B lymphocytes: a requirement for atp-citrate lyase in lipopolysaccharide-induced differentiation. J Biol Chem 289: 7011–7024.

49. Xiao, G., L. N. Chan, L. Klemm, D. Braas, Z. Chen, H. Geng, Q. C. Zhang, A. Aghajanirefah, K. N. Cosgun, T. Sadras, J. Lee, T. Mirzapoiazova, R. Salgia, T. Ernst, A. Hochhaus, H. Jumaa, X. Jiang, D. M. Weinstock, T. G. Graeber, M. and Müschen. 2018. B-cell-specific diversion of glucose carbon utilization reveals a unique vulnerability in B cell malignancies. Cell 173: 470–484.

50. Stincone, A., A. Prigione, T. Cramer, M. M. Wamelink, K. Campbell, E. Cheung, V. Olin-Sandoval, N. M. Grüning, A. Krüger, A. M. Tauqeer, M. A. Keller, M. Breitenbach, K. M. Brindle, J. D. Rabinowitz, and M. Ralser. 2015. The return of metabolism: biochemistry and physiology of the pentose phosphate pathway. Biol Rev Camb Philos Soc 90: 927–963.

51. Averill, R. H., J. Bailey-Serres, and N. J. Kruger. 1998. Co-operation between cytosolic and plastidic oxidative pentose phosphate pathways revealed by 6-phosphogluconate dehydrogenase-deficient genotypes of maize. Plant J 14: 449–457.

52. Koutcher, J. A., A. A. Alfieri, C. Matei, K. L. Meyer, J. C. Street, and D. S. Martin. 1996. Effect of 6-aminonicotinamide on the pentose phosphate pathway: ^31^P NMR and tumor growth delay studies. Magn Reson Med 36: 887–892.

53. Chandel, N. S. 2021. Carbohydrate metabolism. Cold Spring Harb Perspect Biol 13: a040568.

54. Chang, C. H., J. D. Curtis, L. B. Maggi, B. Faubert, A. V. Villarino, D. O’Sullivan, S. C. Huang, G. J. W. van der Windt, J. Blagih, J. Qiu, J. D. Weber, E. J. Pearce, R. G. Jones, and E. L. Pearce. 2013. Posttranscriptional control of T cell effector function by aerobic glycolysis. Cell 153: 1239–1251.

55. Palaskas, N. J., J. D. Garcia, R. Shirazi, D. S. Shin, C. Puig-Saus, D. Braas, A. Ribas, and T. G. Graeber. 2019. Global alteration of T-lymphocyte metabolism by PD-L1 checkpoint involves a block of de novo nucleoside phosphate synthesis. Cell Discovery 5: 1–10.

56. Cocco, M., S. Stephenson, M. A. Care, D. Newton, N. A. Barnes, A. Davison, A. Rawstron, D. R. Westhead, G. M. Doody, and R. M. Tooze. 2012. In vitro generation of long-lived human plasma cells. J Immunol 189: 5773–5785.

57. Panneerselvam, K., J. R. Etchison, and H. H. Freeze. 1997. Human fibroblasts prefer mannose over glucose as a source of mannose for N-glycosylation. Evidence for the functional importance of transported mannose. J Biol Chem 272: 23123–23129.

58. Alton, G., M. Hasilik, R. Niehues, K. Panneerselvam, J. R. Etchison, F. Fana, and H. H. Freeze. 1998. Direct utilization of mannose for mammalian glycoprotein biosynthesis. Glycobiology 8: 285–295.

59. Sharma, V., M. Ichikawa, and H. H. Freeze. 2014. Mannose metabolism: more than meets the eye. Biochem Biophys Res Comm 453: 220–228.

60. Teng, X., J. Brown, S. C. Choi, W. Li, and L. Morel. 2020. Metabolic determinants of lupus pathogenesis. Immunol Rev 295: 167–186.

61. Edwards, D. N., V. M. Ngwa, A. L. Raybuck, S. Wang, Y. Hwang, L. C. Kim, S. H. Cho, Y. Paik, Q. Wang, S. Zhang, H. C. Manning, J. C. Rathmell, R. S. Cook, M. R. Boothby, and J. Chen. 2021. Selective glutamine metabolism inhibition in tumor cells improves antitumor T lymphocyte activity in triple-negative breast cancer. J Clin Invest 131 e140100.

62. Li, H., A. Boulougoura, Y. Endo, and G. C. Tsokos. 2022. Abnormalities of T cells in systemic lupus erythematosus: new insights in pathogenesis and therapeutic strategies. J Autoimmun 21: 102870.

63. Dufort, F. J., M. R. Gumina, N. L. Ta, Y. Tao, S. A. Heyse, D. A. Scott, A. D. Richardson, T. N. Seyfried, and T. C. Chiles. 2014. Glucose-dependent de novo lipogenesis in B lymphocytes: a requirement for atp-citrate lyase in lipopolysaccharide-induced differentiation. J Biol Chem 289: 7011–7024.

64. Jayachandran, N., E. M. Mejia, K. Sheikholeslami, A. A. Sher, S. Hou, G. M. Hatch, and A. J. Marshall. 2018. TAPP adaptors control B cell metabolism by modulating the phosphatidylinositol 3-kinase signaling pathway: a novel regulatory circuit preventing autoimmunity. J Immunol 201: 406–416.

65. Gitlin, A. D., Z. Shulman, and M. C. Nussenzweig. 2014. Clonal selection in the germinal centre by regulated proliferation and hypermutation. Nature 509: 637–640.

66. Mayer, C. T., A. Gazumyan, E. E. Kara, A. D. Gitlin, J. Golijanin, C. Viant, J. Pai, T. Y. Oliveira, Q. Wang, A. Escolano, M. Medina-Ramirez, R. W. Sanders, and M. C. Nussenzweig. 2017. The micro-anatomic segregation of selection by apoptosis in the germinal center. Science 358: eaao2602.

67. Victora, G. D., and M. C. Nussenzweig. 2022. Germinal Centers. Annu Rev Immunol 40: 413–442.

68. Yam-Puc, J. C., L. Zhang, R. A. Maqueda-Alfaro, L. Garcia-Ibanez, Y. Zhang, J. Davies, Y. A. Senis, M. Snaith, and K. M. Toellner. 2021. Enhanced BCR signaling inflicts early plasmablast and germinal center B cell death. iScience 24: 102038.

69. Schoeler, K., B. Jakic, J. Heppke, C. Soratroi, A. Aufschnaiter, N. Hermann-Kleiter, A. Villunger, and V. Labi. 2019. CHK1 dosage in germinal center B cells controls humoral immunity. Cell Death Differ. 26: 2551–2567.

70. Schoeler, K., A. Aufschnaiter, S. Messner, E. Derudder, S. Herzog, A. Villunger, K. Rajewsky, and V. Labi. 2019. TET enzymes control antibody production and shape the mutational landscape in germinal centre B cells.

71. He, M., M. B. Saeed, J. Record, M. Keszei, L. Gonçalves Pinho, L. Vasconcelos-Fontes, R. D’Aulerio, R. Vieira, M. M. S. Oliveira, C. Geyer, L. Bohaumilitzky, M. Thiemann, E. Deordieva, L., Buedts, J. P. Matias Lopes, D. Pershin, L. Hammarström, Y. Xia, X. Zhao, C. Cunningham-Rundles, A. J. Thrasher, S. O. Burns, V. Cotta-de-Almeida, C. Liu, A. Shcherbina, P. Vandenberghe, and L. S. Westerberg. 2022. Overactive WASp in X-linked neutropenia leads to aberrant B-cell division and accelerated plasma cell generation. J Allergy Clin Immunol. 149: 1069–1084.

72. Abbott, R. K., J. H. Lee, S. Menis, P. Skog, M. Rossi, T. Ota, D. W. Kulp, D. Bhullar, O. Kalyuzhniy, C. Havenar-Daughton, W. R. Schief, D. Nemazee, and S. Crott. 2018. Precursor frequency and affinity determine B cell competitive fitness in Germinal Centers, tested with germline-targeting HIV vaccine immunogens. Immunity 48: 133–146.

73. Pratama, A., and C. G. Vinuesa. 2014. Control of TFH cell numbers: why and how? Immunol Cell Biol 92: 40–48.

74. Vinuesa, C. G., M. A. Linterman, D. Yu, and I. C. MacLennan. 2016. Follicular Helper T cells. Annu Rev Immunol 34: 335–368.

75. Wong, K. A., J. A. Harker, A. Dolgoter, N. Marooki, and E. I. Zuniga. 2019. T Cell-Intrinsic IL-6R signaling is required for optimal ICOS expression and viral control during chronic infection. J Immunol 203: 1509–1520.

76. O’Connor, B. P., L. A. Vogel, W. Zhang, W. Loo, D. Shnider, E. F. Lind, M. Ratliff, R. J. Noelle, and L. D. Erickson. 2006. Imprinting the fate of antigen-reactive B cells through the affinity of the B cell receptor. J Immunol 177: 1–21.

77. Paus, D., T. G. Phan, T. D. Chan, S. Gardam, A. Basten, and R. Brink. 2006. Antigen recognition strength regulates the choice between extrafollicular plasma cell and germinal center B cell differentiation. J Exp Med 203: 1081–1091.

78. Kräutler, N. J., D. Suan, D. Butt, K. Bourne, J. R. Hermes, T. D. Chan, C. Sundling, W. Kaplan, P. Schofield, J. Jackson, A. Basten, D. Christ, and R. Brink. 2017. Differentiation of germinal center B cells into plasma cells is initiated by high-affinity antigen and completed by Tfh cells. J Exp Med 214: 1259–1267.

79. Ise, W., and T. Kurosaki. 2019. Plasma cell differentiation during the germinal center reaction. Immunol Rev 288: 64–74.

80. Liu, T., R. J. Kishton, A. N. Macintyre, V. A. Gerriets, H. Xiang, X. Liu, E. D. Abel, D. Rizzieri, J. W. Locasale, and J. C. Rathmell. 2014. Glucose transporter 1-mediated glucose uptake is limiting for B-cell lymphoblastic leukemia anabolic metabolism and resistance to apoptosis. Cell Death & Disease 5: 1–13.

81. Quéméneur, L., L. M. Gerland, M. Flacher, M. Ffrench, J. P. Revillard, and L. Genestier. 2003. Differential control of cell cycle, proliferation, and survival of primary T lymphocytes by purine and pyrimidine nucleotides. J Immunol 170: 4986–4895.

82. Nose, M., and H. Wigzell. 1983. Biological significance of carbohydrate chains on monoclonal antibodies. Proc Natl Acad Sci USA 80: 6632–6636.

83. Gala, F. A., and S. L. Morrison. 2002. The role of constant region carbohydrate in the assembly and secretion of human IgD and IgA1. J Biol Chem 277:29005–29011.

84. Masciarelli, S., A. M. Fra, N. Pengo, M. Bertolotti, S. Cenci, C. Fagioli, D. Ron, L. M. Hendershot, and R. Sitia R. 2010. CHOP-independent apoptosis and pathway-selective induction of the UPR in developing plasma cells. Mol Immunol 47: 1356–1365.

